# Reconstructing whole-organism cell phylogenies with resolved ancestral transcriptional states

**DOI:** 10.64898/2026.07.30.741939

**Authors:** Zhan Liu, Shanjun Deng, Hui Zeng, Mendong Zhang, Boliang Liu, Huiming Xiang, Zhiyun Chen, Ao Zhang, Jay Shendure, Xionglei He

**Affiliations:** State Key Laboratory of Biocontrol, School of Life Sciences, Sun Yat-Sen University, Guangzhou 510275, China; Department of Genome Sciences, University of Washington, Seattle, WA, USA; Howard Hughes Medical Institute, Seattle, WA, USA

## Abstract

Combining cell lineage tracing with single-cell RNA sequencing can reconstruct a phylogenetic tree to integrate single-cell transcriptomic atlas. However, the internal nodes of this tree - representing ancestral cells - remain transcriptionally silent, preventing along-lineage longitudinal tracing of cell state dynamics. Here, we overcome this by reconstructing high-resolution zygote-to-larva developmental cell phylogenies for 15 zebrafish larvae, with directly measured transcriptomes of the terminal nodes (i.e., sampled cells). Leveraging a set of lineage-committed upregulated genes (LUGs), we developed LUG-encoded ancestral projection (LEAP), a novel phylogeny-based computational framework, and successfully imputed the transcriptional states of internal nodes of the phylogenies. This enabled, for the first time in a non-nematode organism, lineage-informed longitudinal analysis of cell state dynamics throughout development. Our analysis revealed a major, previously unappreciated wave of fate specializations associated with hatching, distinct from the well-characterized events of gastrulation. Furthermore, we uncovered abundant incipient cell states that are already fate-determined but exhibit minimal transcriptional differentiation, revealing a hidden layer of developmental fate specializations. In sum, by resolving ancestral transcriptional states of a reconstructed cell phylogeny, this work paves the way for constructing lineage-resolved cell atlases in complex organisms to characterize comprehensive cell state dynamics.

## Introduction

The emerging capacity of reconstructing cell phylogenetic trees in complex organisms to understand lineage dynamics and cell state transitions transformed the studies on development and disease progression in the past decade (*1–9*). The cutting-edge methods are now able to reconstruct developmental cell phylogenies, with measured transcriptomes available for the terminal nodes (i.e., the sampled cells) (*10–15*). Nevertheless, from an experimental perspective, accurate and scalable whole-organism cell phylogeny reconstruction remains technically challenging, and only a limited number of systems have achieved sufficient resolution and coverage for organism-scale lineage analysis(*16–18*). Furthermore, the internal nodes, representing the past ancestors of the sampled cells, have no molecular information available because they are not sampled(*19–21*). The lack of ancestral transcriptional states in a reconstructed cell phylogeny prevents true longitudinal tracing of cell-state dynamics, posing a major challenge to the field (*22–25*).

In this study, we addressed this challenge by reconstructing high-resolution cell phylogenies with resolved ancestral transcriptional states in zebrafish, thereby enabling lineage-informed longitudinal analysis of cell-state dynamics in a non-nematode organism. Specifically, we applied SMALT, a powerful cell lineage tracing system developed in our laboratory(*16*, *18*, *26*), to zebrafish and obtained, in 15 zebrafish larvae (7 dpf), high-resolution zygote-to-larva developmental cell phylogenies, with transcriptomes of the terminal nodes directly measured. From these transcriptomes, we characterized a set of lineage-committed upregulated genes (LUGs), which leave expression signatures at the internal nodes. Using *C. elegans* as a benchmark, we then developed LUG-encoded ancestral projection (LEAP), a phylogeny-based computational framework that leverages single-cell transcriptomes from matched developmental stages as references to impute transcriptional states and developmental stages of internal nodes in the zebrafish cell phylogenies. This enabled extensive lineage-informed longitudinal tracing of cell state dynamics throughout development, revealing a global map of fate specializations with novel insights into vertebrate development.

## Results

### Establishment of scSMALT in zebrafish

The SMALT system in this study comprises a base editor of three functional domains and two separate DNA barcodes for lineage recording (Fig. 1a). Among the three domains of the base editor, the AID domain is responsible for catalyzing cytosine-to-uracil transitions, iSceI for DNA binding, and UGI for suppressing the repair of uracil(27–29). The two barcode sequences (HMF1 and HMF2) are each engineered to be one kilobase in size, featuring secondary structures optimized for AID mutagenesis, and harbor 16 iSceI binding sites (each 18 base pairs long and not found in the zebrafish genome). Consequently, the base editor can selectively bind the barcodes to continuously induce C·G-to-T·A substitutions during development. To facilitate integration with single-cell RNA sequencing (scRNA-seq), the two barcodes are each designed to be within the 3’-UTR of a ubiquitously expressed GFP (Fig. 1a).

**Figure 1.**
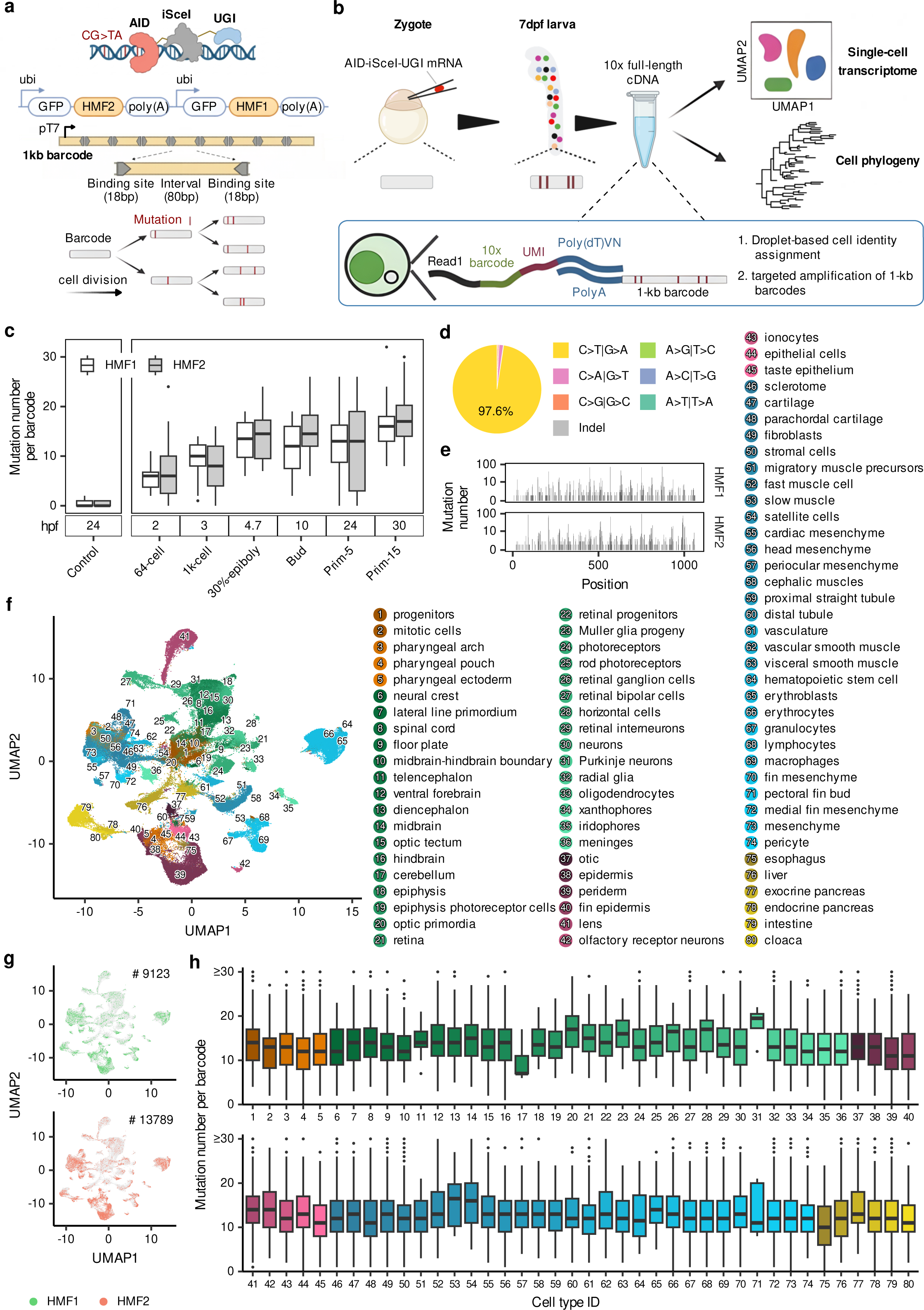
Single-cell RNA-integrated SMALT (scSMALT) in zebrafish. **a.** Schematic overview of the scSMALT system. The base editor, consisting of AID, iSceI, and UGI domains, targets two 1-kb barcode cassettes. Mutations accumulate during cell division and serve as heritable lineage records, as shown below. **b.** Experimental workflow for scSMALT in zebrafish. The procedure involves microinjection of AID-iSceI-UGI mRNA into embryos, followed by the co-capture of single-cell transcriptomes and lineage barcodes during library construction. **c.** Temporal accumulation of mutations during zebrafish development. Boxplots displaying the number of mutations per barcode across developmental stages, ranging from 2 hpf to 30 hpf. Control samples representing embryos without AID-iSceI-UGI mRNA injection were shown on the left. **d.** Mutational spectrum of the scSMALT system. Pie chart illustrating the distribution of base substitution types observed in genomic DNA, with segments colored according to the specific mutation class. **e.** Positional distribution of mutations along the 1-kb barcodes. Histograms illustrating the mutation frequency at each nucleotide position within the HMF1 and HMF2 barcode cassettes. **f.** UMAP projection of 80 annotated cell types at 7 dpf. Cells are color-coded by identity, with color palettes grouped to reflect major developmental compartments. **g.** Recovery of lineage barcodes across the cell population. UMAP plots highlighting cells with recovered HMF1 and HMF2 lineage barcodes. **h.** Mutation counts across cell-type identities. Boxplots show the number of mutations per barcode for each of the 80 cell types for HMF1 and HMF2. The x-axis indicates cell-type IDs, corresponding to the numerical annotations defined in panel f.

As such, single-cell RNA sequencing can simultaneously capture the endogenous transcripts and the barcode transcript of a cell (Fig. 1b). In addition, a T7 promoter was inserted between the GFP gene and the 1-kb barcode, allowing specific amplification of the 1-kb barcode from cDNA libraries (Fig. 1a; Methods). Collectively, these components established single-cell RNA-integrated SMALT (scSMALT) in zebrafish. We generated a transgenic zebrafish line carrying the dual-barcode cassette (Fig. S1; Methods) and induced mutagenesis by injecting AID-iSceI-UGI mRNA at the one-cell stage. While negligible mutations were observed in control embryos, the lineage barcodes from genomic DNA in the mRNA-injected group accumulated mutations progressively throughout development, despite substantial variations across individuals (Fig. 1c). Specifically, the mean mutation count of HMF1 increased to 11.4 by the end of gastrulation (10 hours post fertilization, 10 hpf) and continuously accumulated to 16.2 at 30 hpf, a trend consistently mirrored by HMF2 (Fig. 1c). This accumulation rate, approximating one mutation per cell division (Fig. S2; Methods), suggests a high-resolution barcoding of the cell divisions. Consistent with the AID-induced mutation profile, the barcoding mutations were predominantly (97.7%) C·G-to-T·A substitutions (Fig. 1d). In addition, the mutations were distributed throughout each barcode sequence, highlighting the large state space of potential barcodes (Fig. 1e).

We then applied scSMALT to zebrafish and obtained 10x Genomics scRNA-seq data for 15 larvae (7 days post fertilization, dpf; Methods). Our dataset encompasses 165,953 high-quality single cells, which were classified into 80 major cell types via unsupervised clustering and manual annotation(*30–33*) (Fig. 1f). Target amplification from the 10x Genomics cDNA libraries followed by PacBio HiFi sequencing successfully recovered HMF1 and HMF2 in 9,123 and 13,789 cells, respectively (Fig. 1g; Fig. S3; Methods). Mutations were then identified for the recovered barcodes of each cell (Methods). There were, on average, 14.6 mutations in HMF1 and 11.7 in HMF2 (Fig. S4). The slightly lower mutation count in HMF2 reflected a recurrent depletion in cDNA amplicons of a ∼300bp region of the barcode (Fig. S5; Methods). Both barcodes were recovered in cells of all 80 annotated cell types despite variations in abundance (Fig. S6), and the average mutation counts exhibited slight fluctuations across the cell types (Fig. 1h). In addition, the vast majority of mutations (93.1%) were C·G-to-T·A substitutions (Fig. S7a), with wide distribution along the barcodes (Fig. S7b). Within each individual, barcode diversity at the terminal level was high, with an average of 90.9% of the recovered 1-kb barcodes being unique (Fig. S8). Together, these results established scSMALT as a promising system for resolving the developmental lineage history of zebrafish.

### Zygote-to-larva developmental cell phylogenies reconstructed by scSMALT

Using a maximum-likelihood method(*34*, *35*), we conducted phylogeny reconstruction based on the barcoding mutations in HMF1 and HMF2, respectively, for each of the fifteen 7 dpf larvae (Methods). This resulted in a total of 30 zygote-to-larva developmental cell phylogenies (Fig. 2a-b). On average, each cell phylogeny comprised 764 terminal nodes/cells (cells sampled at 7 dpf), ranging from 134 to 1,648 (Fig. 2c). The transcriptomes of the terminal cells were then assigned according to the 10x Genomics cell barcode sequence attached with each barcode (Fig. 2a-b; Methods).

**Figure 2.**
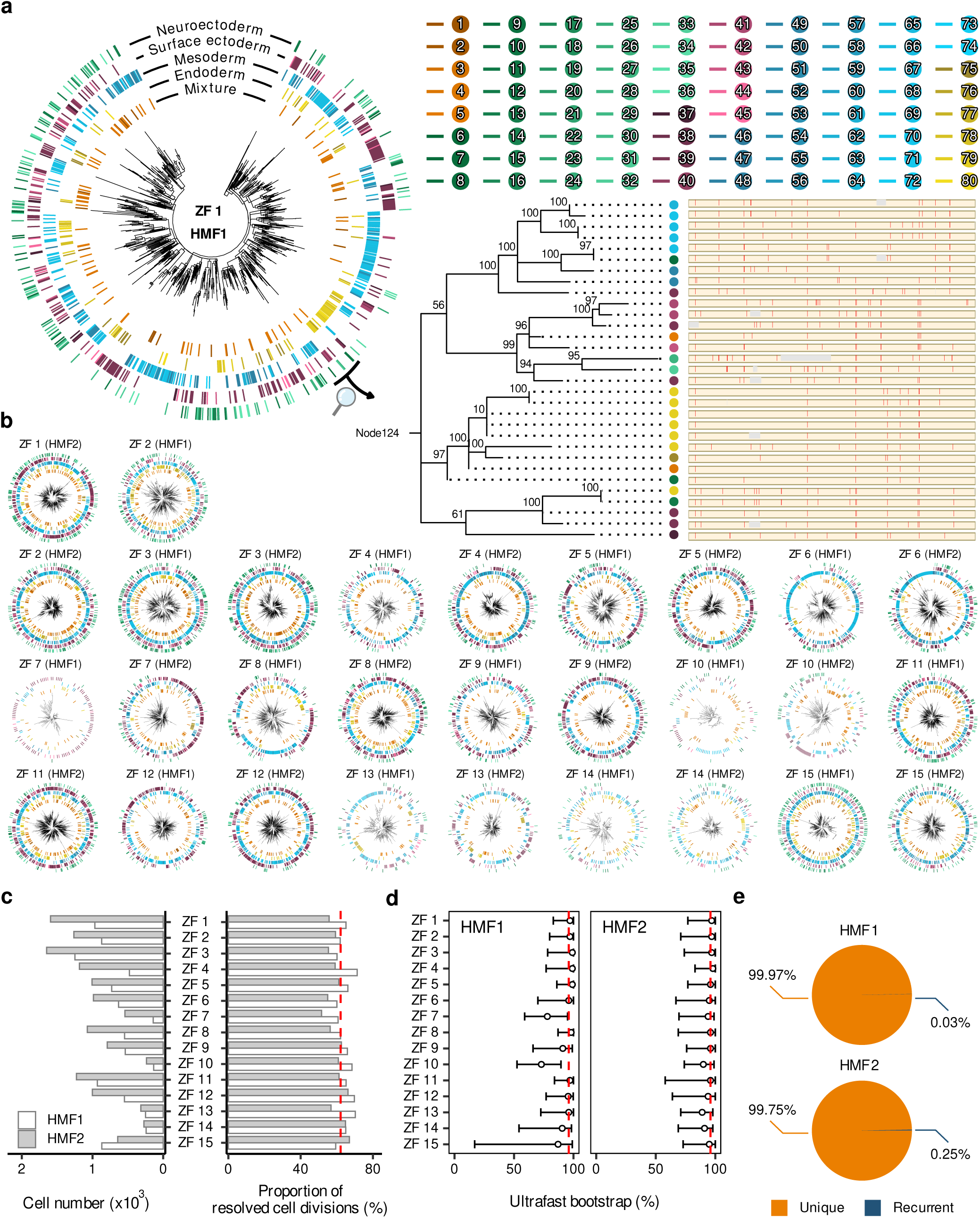
High-resolution zygote-to-larva developmental cell phylogenies reconstructed by scSMALT. **a.** Representative cell phylogeny derived from scSMALT data. The outer circular tracks annotate major developmental compartments and 80 terminal cell-type identities. The magnified subtree (right) highlights the ultrafast bootstrap values at internal nodes alongside the barcode mutation matrix for terminal cells. **b.** Maximum-likelihood cell phylogenies across 15 individuals. Reconstructed phylogenetic trees for both HMF1 and HMF2 lineage barcodes are presented for individuals ZF1–ZF15, following the visualization format described in panel a. **c.** Statistics of terminal cell number and phylogenetic resolution. Bar plots show the total number of terminal cells (left) and the proportion of resolved internal nodes (right) across all 30 reconstructed phylogenies. The dashed red line denotes the mean proportion of resolved internal nodes. **d.** Distributions of ultrafast bootstrap values for internal nodes. Boxplots illustrating support values for HMF1 and HMF2 phylogenies across the 15 individuals. The dashed red line denotes the mean ultrafast bootstrap value across the 15 corresponding phylogenies. **e.** Barcode recurrence across individuals. Pie charts illustrating the proportions of unique versus recurrent lineage barcodes observed for HMF1 and HMF2 across all 15 individuals.

The 30 reconstructed cell phylogenies were generally of high quality. Overall, about 62.1% of ancestral cell division events were resolved, corresponding to 13,948 internal nodes across the 30 phylogenies (Fig. 2c; Methods). These internal nodes were statistically supported by high ultrafast bootstrap values, with a median of 96% for both HMF1- and HMF2-based phylogenies (Fig. 2d). The resolving rate was reduced near the root, likely reflecting rapid cell divisions at cleavage stages (Fig. S9). To assess potential confounding effects from barcode homoplasy, we examined identical barcodes across individuals. Such identical barcodes were found rare (0.03% for HMF1 and 0.25% for HMF2; Fig. 2e), indicating minimal barcode homoplasy. Together, the 30 cell phylogenies were statistically reliable, with high resolution in resolving the lineage history of the sampled cells.

We further checked how phylogenetic positions explain transcriptomes of the sampled cells. Overall, cells of the same type in the transcriptome tended to be phylogenetically clustered (Fig. 2a; Fig. S10; Methods), indicating a correspondence between lineage proximity and transcriptional similarity. The transcriptional similarity was quite high between sister nodes of a phylogeny, but decayed rapidly with increasing genealogical distance, approaching background levels after three to four branching events (Fig. S11). The rapid decay was consistent with the observation in *C. elegans*(*36*). Beyond this global trend, we observed substantial heterogeneity in the phylogeny-coupled transcriptional divergence across cell types and germ layers (Fig. S12), suggesting that lineage-dependent constraints on transcriptional programs are dynamically regulated during zebrafish development.

### LEAP as a phylogeny-based method for imputing ancestral transcriptional states

The inference of ancestral cell states of a reconstructed cell phylogeny can be achieved by discovering lineage-coupled molecular features(*25*). During cell fate commitment, some genes become activated or silenced at certain time points and then remain the state in all descendant lineages; such lineage-committed expression signatures would enable the inference of ancestral states from terminal states (Fig. 3a). As in our previous study(*25*), the expression levels of a gene across different cells can be binarized as upregulated (expression state = 1) and downregulated (expression state = 0) to approximate activation and silence in analyzing single-cell transcriptomes; because the high dropout rate in single-cell transcriptomes confounds the detection of downregulation, we focused on lineage-committed upregulated genes (LUGs) (Methods). For an LUG that shows upregulation in the terminal cells of a given phylogenetic clade, the internal nodes of the clade can all be assigned the same upregulation state for this LUG (Fig. 3a). Combining the states of many LUGs results in an LUG-encoded transcriptional profile for each internal node, which can then be projected onto a reference transcriptomic atlas to infer the cell identity of the internal nodes. Based on this reasoning, we developed LEAP (LUG-encoded Ancestral Projection), a computational framework for imputing ancestral transcriptional states of a reconstructed cell phylogeny (Fig. S13; Methods).

**Figure 3.**
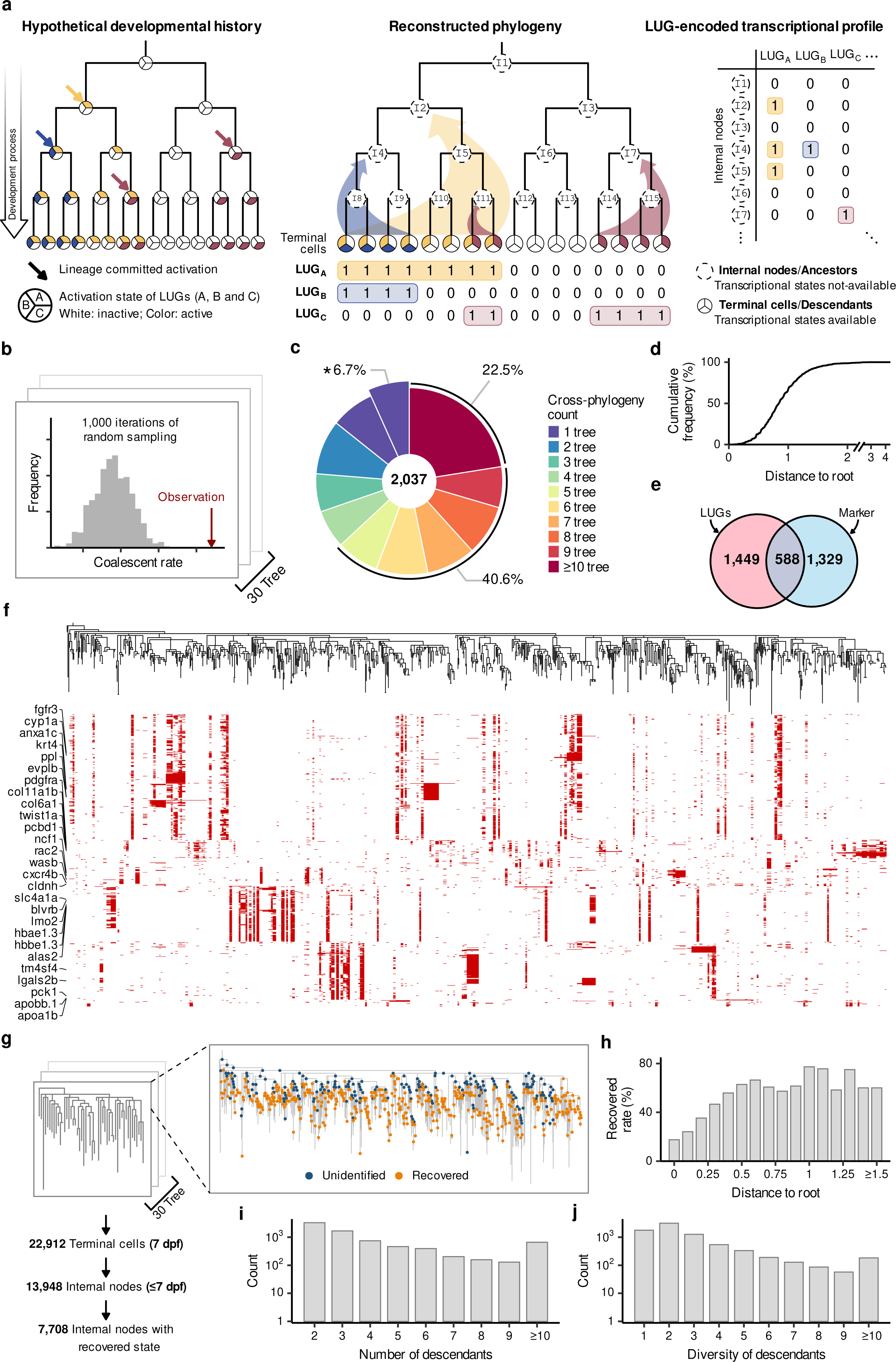
LUG-encoded reconstruction of ancestral transcriptional states. **a.** Schematic of the LEAP framework for ancestral state projection. The framework integrates a hypothetical developmental history (left), defines lineage-committed upregulated genes (LUGs) within the reconstructed phylogeny (middle), and reconstructs ancestral transcriptional states (right). **b.** Statistical definition of LUGs. The null distribution is generated through 1,000 iterations of random sampling; the observed value is marked by the red arrow. **c.** Reproducibility of LUGs across phylogenies. Donut chart displaying the distribution of 2,037 LUGs by their recurrence across 30 cell phylogenies. The asterisk marks LUGs identified in only one phylogeny. **d.** Cumulative frequency of LUG activation nodes. Distribution of inferred LUG activation events plotted as a function of phylogenetic distance from the root. **e.** Overlap between LUGs and cell-type markers. Venn diagram illustrating the intersection between the identified LUGs and known cell-type markers. **f.** Distribution of LUGs across a representative cell phylogeny. Each red tile represents a LUG upregulated in the corresponding phylogenetic clade. Well-characterized development relevant genes are labeled on the left. **g.** Summary of ancestral state recovery across 30 phylogenies. Flowchart illustrating the workflow and data quantities, tracking the number of terminal cells, reconstructed internal nodes, and the subset of internal nodes with successfully recovered LUG-encoded transcriptional states. **h.** Recovery rate of ancestral states. Recovery rate of LUG-encoded internal node states plotted as a function of phylogenetic distance from the root. **i.** Distribution of descendant cell counts. Histogram showing the number of terminal descendants associated with each internal node where a transcriptional state was recovered. **j.** Diversity of descendant cell types. Histogram showing the number of distinct terminal cell types derived from each internal node where a transcriptional state was recovered.

To benchmark LEAP before applying it to zebrafish, we leveraged the well-studied cell phylogenetic tree of *C. elegans*, accompanied by its lineage-resolved transcriptomic atlas(*36*), as a biological ground truth. Using a previously developed algorithm TarCA(*25*), we first identified 4,788 LUGs that exhibit clade-restricted upregulation patterns in the phylogeny (Fig. S14a; Methods). The LUG-encoded transcriptional profiles of the internal nodes were then co-embedded with the reference transcriptomic atlas, revealing strong concordance in UMAP space with the local inverse Simpson’s Index (iLISI)(*37*) score being 1.44 and 1.62 before and after batch correction, respectively (Fig. S14b-c; Methods). We therefore performed label transfer from the reference atlas to the internal nodes based on the LUG-encoded transcriptional profiles (Methods). We achieved generally high accuracy in cell identity assignments (Fig. S14d). For instance, the lineage assignment accuracy was 100%, 100%, 86.7%, 90.9%, and 51.6% for the five founder lineages AB, C, D, E, and MS, respectively (Fig. S14e). At single-cell resolution, 47% of the internal nodes were assigned to their exact identities, representing nearly a ten-fold enrichment over background (Fig. S14f). This accuracy is not low, given the fact that many cells’ transcriptomes are nearly indistinguishable in the nematode(*36*). As a matter of fact, the assignment accuracy for cell-type categories consistently exceeded 90% (Fig. S14g). These results collectively justified LEAP as a promising tool.

We next applied LEAP to the zebrafish cell phylogenies. Using TarCA, we identified 2,037 LUGs (Supplementary Table 4), 85.3% of which were detected in at least two phylogenies and 63.1% in five or more (Fig. 3c; Methods). The activation timing of LUGs is distributed across various phylogenetic depths (Fig. 3d), consistent with the progressive emergence of lineage-committed expression programs. Notably, LUGs were enriched not only for terminal cell-type markers (Fig. 3e), but also for early germ layer-associated genes (Fig. S15). As in an exemplar phylogeny shown in Fig. 3f, hematopoietic regulators (*lmo2*)(*38*) and mesenchymal genes (*pdgfra*, *twist1a*)(*39*, *40*) were confined to distinct mesodermal clades; meanwhile, epithelial markers (*krt4*, *ppl*)(*41*, *42*) and endodermal regulators (*apoa1b*, *apobb.1*)(*43*, *44*) were highly enriched within their respective, non-overlapping lineage clades. These patterns aligned with their established roles in fate specification, validating that LUGs capture genuine lineage-committed programs.

As in *C. elegans*, we constructed the LUG-encoded transcriptional profiles for the 13,948 internal nodes of the 30 zebrafish cell phylogenies (Fig. 3g). A total of 7,708 internal nodes contain an informative profile with at least three LUGs being activated (expression state = 1). The rate of recovering an informative profile increased from ∼ 20% for internal nodes near the root to ∼70% for those distant from the root (Fig. 3h), consistent with the reduced phylogenetic resolution at early stages. The recovered internal nodes each seeded an average of 4.6 terminal cells and 2.8 cell types (Fig. 3i-j), highlighting the multipotency of these internal nodes.

### Successful inferences of ancestral states of the zebrafish cell phylogenies

To assign cell states of the internal nodes, we projected the LUG-encoded transcriptional profiles onto a time-resolved zebrafish scRNA-seq reference atlas(*30*). This atlas comprises nearly half a million cells spanning 136 cell types across 14 stages from 3hpf to 120 hpf, providing a broad coverage of developmental cell states. Co-embedding of the internal nodes with the reference atlas revealed very strong concordance in UMAP space, with an iLISI score of 1.79 and 1.81 before and after batch correction, respectively (Fig. 4a; Fig. S16; Fig. S17). Each internal node was then mapped to its nearest neighbors of the reference atlas, enabling joint assignment of cell-type identity and developmental stage (Methods).

**Figure 4.**
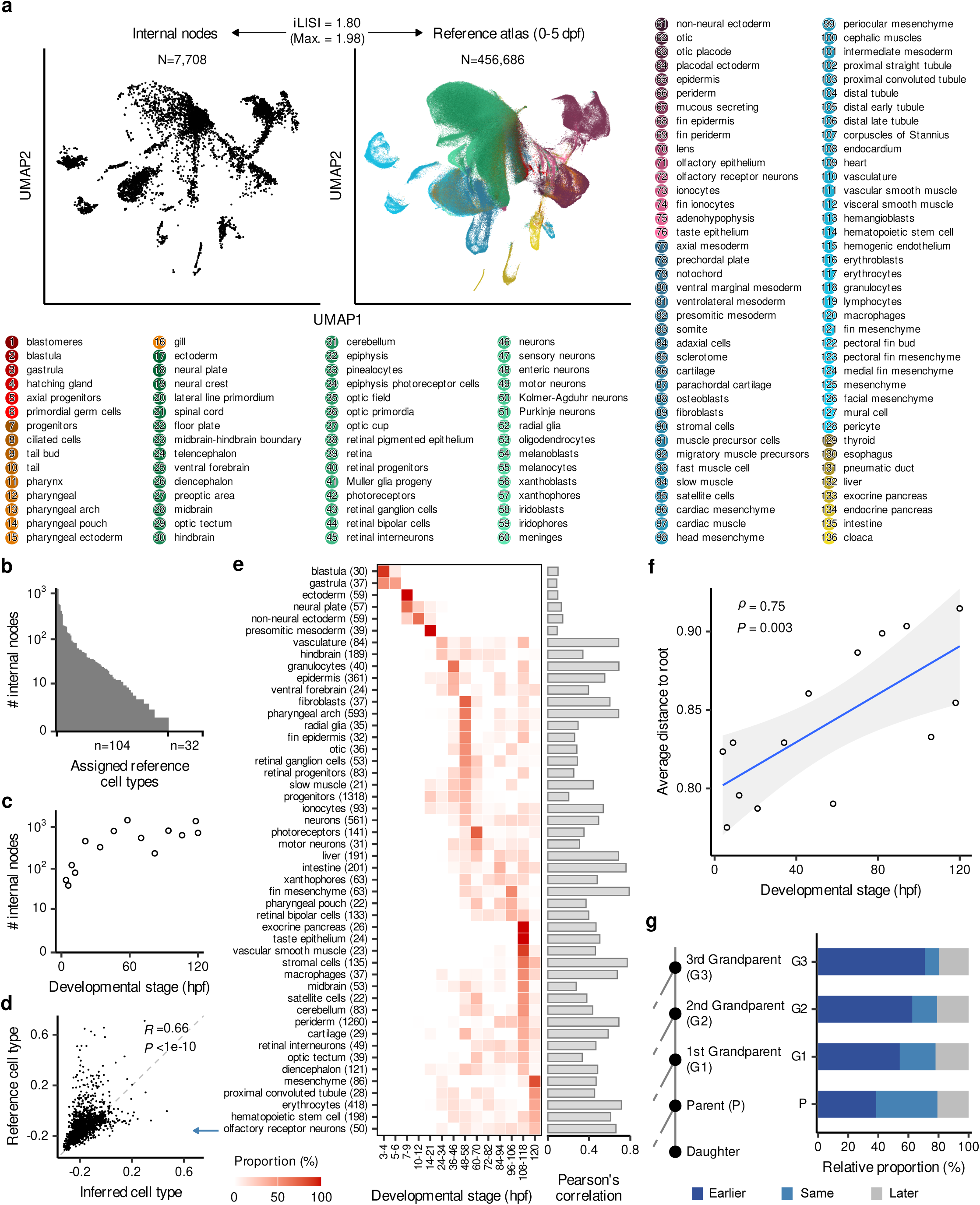
LEAP enables accurate imputation of ancestral states in zebrafish phylogenies. **a.** UMAP co-embedding of inferred internal nodes and a reference cell atlas. UMAP visualization showing the CCA integration of 7,708 inferred internal nodes (left) with 456,686 reference cells (right). Reference cells are color-coded by the 136 annotated cell types listed below. The iLISI score are attached on the top. **b.** Distribution of mapped ancestral cell types. Histogram showing the assignment of inferred internal nodes across the 136 reference cell types, with 104 types successfully recovered and 32 types absent. **c.** Temporal distribution of internal nodes. Scatter plot showing the number of inferred internal nodes across developmental stages, ranging from 3 to 120 hpf. **d.** Transcriptional correlation between inferred and reference cell types. Scatter plot comparing scaled expression profiles of reconstructed internal nodes against reference cells for olfactory receptor neurons. The Pearson correlation coefficient and P-value are attached. **e.** Global characterization of inferred cell types. The left panel shows the proportional distribution of mapped internal nodes across developmental stages (3–120 hpf) for 39 cell types with >20 cases. The right panel displays the corresponding Pearson’s correlation coefficients between reconstructed and reference transcriptional profiles. **f.** Correlation between phylogenetic depth and inferred developmental timing. Scatter plot of the average distance to the root (y-axis) versus inferred developmental stage in hpf (x-axis). The Spearman correlation coefficient and P-value are attached. **g.** Consistency of inferred developmental stages with phylogenetic positions. For internal nodes along the same lineage, parental nodes are often mapped to earlier or the same developmental stages relative to the daughter.

A total of 104 annotated cell types were successfully assigned to the internal nodes (Fig. 4b-c), encompassing developmental stages from 3 hpf to 120 hpf (Fig. 4c). To assess the assignment fidelity, for each cell type, we compared the aggregated LUG-encoded transcriptional profiles of the internal nodes against the corresponding reference cells. Across the 39 cell types with over 20 assigned internal nodes, we observed quite robust transcriptional concordance (Fig. 4d-e; Fig. S18; Methods). The correlations were modestly reduced in early-stage populations, including blastula and gastrula cells, which is explained by limited cell number and phylogenetic resolution near the root (Fig. 4e). Nonetheless, the generally high correlations for the remaining cell types indicated the success of LEAP in imputing transcriptional states of the internal nodes.

Beyond cell-type identity, a high-fidelity label assignment should adhere to the inherent temporal logic of embryogenesis: deeper internal nodes in a phylogeny, representing earlier ancestors, should map to earlier developmental stages. To test this, we checked the overall relationship between phylogenetic depth and inferred developmental stage of the internal nodes, and a robust positive correlation was observed (Spearman’s ρ = 0.75, P = 0.003; Fig. 4f). For internal nodes along the same lineage, ∼80% of parental nodes were assigned to earlier or the same developmental stages compared to their descendants, and this temporal congruency strengthened with increasing lineage distance (e.g., from parents to grandparents; Fig. 4g; Fig. S19), highlighting the performance of LEAP

To further assess the ancestral inferences from a functional perspective, we performed perturbation-based single-cell phenotyping. We focused on the imputed cell population designated as ‘neurons’ and selected the top five ranked LUGs (*dnajc5aa*, *smarcd1*, *tra2b*, *fam168b*, and *pdcl*) identified from ‘neurons’ (Fig. 5a-b; Methods). Because LEAP assigned the highest proportion of these ‘neurons’ to the 48–58 hpf window (Fig. S20), we focused our phenotypic assessments on embryos at 48hpf. Specifically, we employed scRNA-seq to profile both wild-type (WT) embryos and F0 knockouts (crispants) (*45*) generated via CRISPR–Cas9 mutagenesis (Fig. 5c; Fig. S21; Methods). In total, we captured 38,125 high-quality cells from 18 individual WT embryos to establish a baseline reference, alongside 136,615 cells from 53 F0 crispants (8–12 individuals per perturbation) (Fig. 5c; Fig. S22; Methods). To visualize global compositional shifts, we performed dimensionality reduction on normalized embryo-level cell-type abundances (Methods)(*46*). The resulting UMAP embedding showed that embryos clustered predominantly by genotype, indicating that specific genetic perturbations were the primary drivers of cellular variance (Fig. 5e; Methods). Compared to the wild-type baseline, we observed statistically significant reductions of ‘neurons’ in all five knockout genotypes, with an average of 72.3% drop in normalized cell counts (Fig. 5f; Methods). Notably, similar reductions were observed in other neuroectoderm-derived cell types (Fig. S23), suggesting the broad neuronal functions of these LUGs.

**Figure 5.**
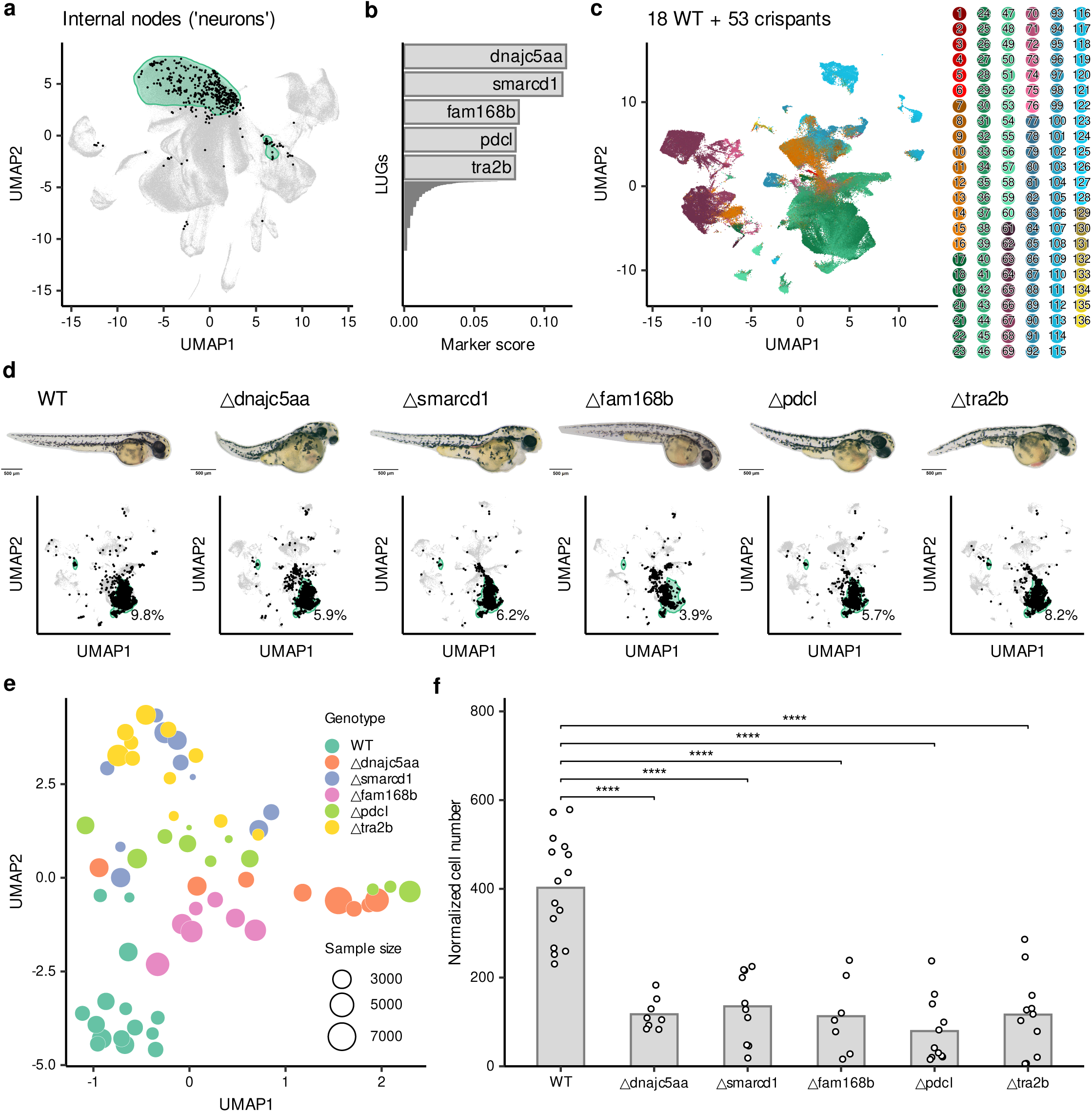
Perturbation-based functional assessment of ancestral inferences. **a.** UMAP embedding highlighting internal nodes (black) assigned to ‘neurons’ based on their LUG-encoded transcriptional profiles. The shadow in green highlights the ’neurons’ population in reference atlas. **b.** Top ‘neurons’-enriched LUGs. Bar plot displaying marker scores for the top five candidates. **c.** Integrated single-cell transcriptomes. UMAP visualization of 18 wild-type (WT) and 53 F0 crispant embryos at 2 dpf, with cells color-coded by annotated cell type. **d.** Phenotypic and transcriptomic assessment of crispants. Representative brightfield images of WT and F0 crispant embryos at 2 dpf (top) and corresponding UMAP embeddings (bottom) highlighting the ‘neurons’ population (shadow in green). Percentages indicate the relative abundance of ‘neurons’ in each genotype. **e.** Embryo-level cell-type abundance. UMAP of normalized cell-type compositions. Dot color represents genotype, and dot size reflects the number of cells recovered per embryo. **f.** Normalized cell numbers of ‘neurons’. Bar plot comparing ‘neurons’ abundance between WT and crispant groups. Individual dots represent biological replicates (embryos). Statistical significance was assessed using a Wilcoson test; ’****’ denotes P<0.0001.

### Phylogeny-informed longitudinal analysis reveals a major wave of fate specializations during the embryo-to-larva transition

Establishing the transcriptional states of internal nodes enabled a lineage-informed longitudinal analysis of cell-state dynamics. For internal nodes with assigned cell-type identities and developmental stages, the potency of differentiating into different terminal fates is not directly available but can be estimated within a phylogeny (Fig. 6a). Conceptually, an internal node giving rise to a single terminal cell type is unipotent, whereas internal nodes seeding diverse cell types are multipotent (Fig. 6b). Quantitatively, the potency of an internal node can be measured using the Shannon entropy of its terminal cell-type composition (Methods).

**Figure 6.**
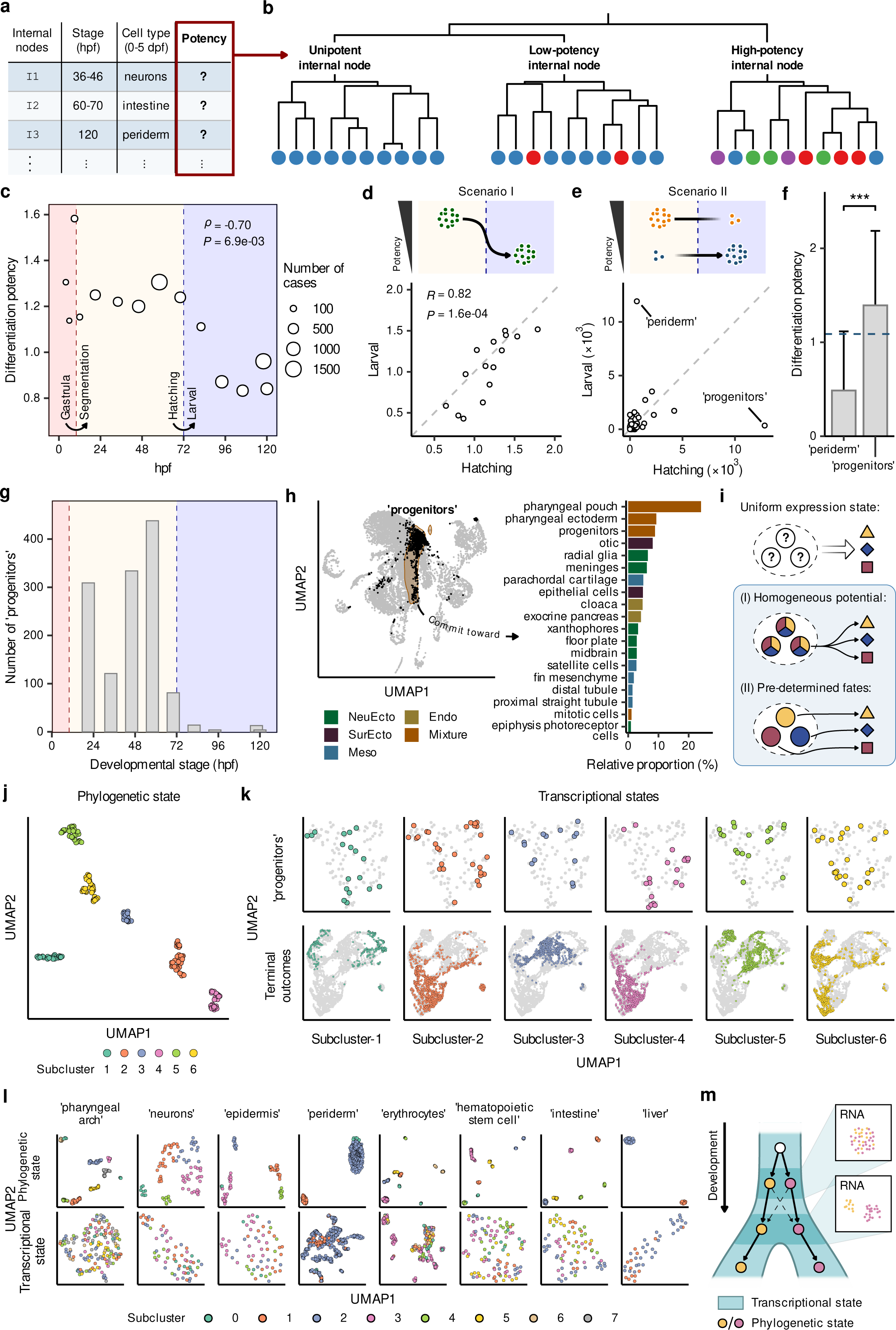
Hatching-associated fate specialization and incipient cell fate commitments. **a.** Schematic representation of internal node characterization. **b.** Conceptual framework for internal-node potency. Internal nodes giving rise to a single terminal cell type are defined as unipotent (left). Nodes contributing to limited or broad sets of terminal descendant cell types are classified as low-potency (middle) and high-potency (right), respectively. **c.** Differentiation potency across zebrafish development. Each point represents internal nodes grouped by developmental time, with point size indicating the number of cases. Spearman’s correlation coefficient and P value are shown. Dashed lines mark the gastrula-to-segmentation and hatching-to-larval transitions. **d.** Scenario I: potency reduction within established cell types. The schematic illustrates a model in which differentiation potency decreases within the same cell types during the hatching-to-larval transition. The scatter plot compares cell-type-specific potency between the hatching and larval periods. **e.** Scenario II: population-level compositional turnover. The schematic illustrates a model in which the overall decrease in potency is driven by contraction of multipotent populations and expansion of fate-restricted populations. The scatter plot compares cell-type abundance between the hatching and larval periods, highlighting the ‘periderm’ and ‘progenitors’ populations. **f.** Differentiation potency of ‘periderm’ and ‘progenitors’. Bar plots show mean differentiation potency with error bars. Statistical significance is indicated; ***P < 0.001. The dashed line indicates the mean potency across all internal nodes. **g.** Temporal distribution of ‘progenitors’. The histogram shows the number of internal nodes assigned to ‘progenitors’ across developmental time. **h.** Transcriptional identity and terminal fate of ‘progenitors’. UMAP visualization highlighting the ‘progenitors’ population within the LUG-encoded transcriptional state space (left). Relative proportions of terminal cell types derived from ‘progenitors’, color-coded by major developmental compartment (right). **i.** Conceptual models of incipient cell fate commitment. Schematic representation of a uniform transcriptional state, illustrating two possible scenarios: (I) homogeneous developmental potential, or (II) predetermined fates that are not yet distinguishable within the transcriptional state space. **j.** Phylogenetic-state clustering of ‘progenitors’. UMAP visualization of the ‘progenitors’ population, color-coded by six subclusters identified based on their terminal descendant cell-type composition. **k.** Correspondence between phylogenetic and transcriptional states. Top row: ‘progenitors’ internal nodes color-coded by the six phylogenetic states identified in (j). Bottom row: Terminal descendant cells for each ‘progenitors’ subcluster projected onto the transcriptional space. **l.** Phylogenetic-state and transcriptional-state relationships across representative cell types. For each cell type, the top row shows internal nodes grouped by phylogenetic state, whereas the bottom row shows the same nodes projected into the LUG-based transcriptional state space and colored by phylogenetic-state subclusters. **m.** Decoupling of phylogenetic and transcriptional states. Differentiation involves an early resolution of cell fate at the phylogenetic level, which temporally precedes the divergence of cellular transcriptional states.

We applied this framework to map global potency dynamics along zebrafish development from 3 to 120 hpf (Fig. 6c). Overall, cell potency exhibited a progressive decline over time (Spearman’s ρ = -0.70, P = 0.0069; Fig. 6c), consistent with the expected pattern of fate specializations during development(*31*, *33*). Comparing cell potency between early developmental stages (up to the end of gastrulation, <10 hpf) and later embryonic periods (10–72 hpf) revealed a discernible decline, aligning with the extensive fate specializations associated with gastrulation (Fig. S24). Notably, the limited number of early internal nodes recovered in the phylogenies may have blurred the comparison (Fig. 6c). Rather than declining continuously, observed potency then stabilized into a prolonged plateau during organogenesis, encompassing the segmentation, pharyngula, and hatching periods (10-72 hpf; Fig. 6c). However, this relative stability was followed by a pronounced reduction post-hatching (>72 hpf; Fig. 6c), reflecting a second, previously unappreciated, major wave of fate specializations. This later wave temporally coincides with the embryo-to-larva transition(*47*).

The hatching-associated decline of potency may be ascribed to two factors: an intrinsic reduction of potency within established cell types (Scenario I) or compositional turnover of the cell types with distinct potency levels (Scenario II) (Fig. 6d-e). Systematically comparing cell-type-specific potency between the hatching and early larval periods revealed that intrinsic potency metrics remained relatively stable for most cell types (Fig. 6d). Instead, we observed a pronounced compositional shift, characterized by the contraction of a multipotent ‘progenitors’ population and the expansion of a fate-restricted ‘periderm’ population (Fig. 6e-f). In silico exclusion of these two populations largely erased this wave of potency decline (Fig. S25), confirming the contribution of the two cell types. The expansion of the ‘periderm’ population after hatching is consistent with continued larval growth and its role as the outer epithelial layer, whereas the factors driving the reduction of the multipotent ‘progenitors’ population remain less clear. We therefore focused on the ‘progenitors’ in the following analysis.

### Incipient fate commitments informed by cell phylogenetic states

The ‘progenitors’ population, as an imputed ancestral cell type, was predominantly present from the segmentation to hatching stages (10–72 hpf) and became largely absent after hatching, a pattern revealed by both phylogenetic and transcriptomic analyses (Fig. 6g; Fig. S26). Despite the fact that major germ layers are established before segmentation, examination of the cell phylogenies showed that this ‘progenitors’ population seeded a wide range of cell types across multiple germ layers (Fig. 6h; Fig. S27; Methods).

We reasoned that the ‘progenitors’ may represent either a homogeneous multipotent population or a heterogeneous pool with diverse hidden fates (Fig. 6i). This can be informed by analyzing the terminal cell-type composition for each internal node labeled as ‘progenitors’ in the cell phylogenies (Methods). Remarkably, clustering based on the terminal cell-type composition revealed six discrete clusters for this ‘progenitors’ population (Fig. 6j; Fig. S28; Methods), each exhibiting distinct biases towards specific terminal cell types (Fig. S29). This suggested that the ‘progenitors’ population, despite being transcriptionally similar, is a heterogeneous pool with already-determined fate biases. These subclusters, representing phylogenetic states defined by terminal-fate composition within a cell phylogeny, likely correspond to incipient fate commitments when transcriptional difference is minimal but fates are already determined. Consistently, despite being clearly partitioned into distinct phylogenetic states, these ‘progenitors’ exhibited highly similar LUG-encoded transcriptional profiles, with extensive overlaps within the transcriptional manifold (Fig. 6k, top). While these ancestral ‘progenitors’ were transcriptionally similar, they ultimately gave rise to distinct terminal cells with divergent transcriptomes (Fig. 6k, bottom).

We extended this analysis to other imputed ancestral cell types to search for more incipient fate commitments. For the eight cell types with over 30 internal nodes assigned, including ‘epidermis’, ‘erythrocytes’, ‘hematopoietic stem cell’, ‘intestine’, ‘neurons’, ‘periderm’, ‘liver’, and ‘pharyngeal arch’, discrete cell phylogenetic states were consistently observed (Fig. 6l; Fig. S30S). In contrast, the corresponding transcriptomic representations were either weakly structured or extensively mixed (Fig. 6l). Hence, our analyses suggested abundant incipient fate commitments that exhibit minimal transcriptional divergences, highlighting the strength of phylogenetic analysis in informing fate determination prior to transcriptional analysis (Fig. 6m).

## Discussion

The field has witnessed growing efforts to reconstruct cell phylogenies to inform transcriptional dynamics during development or disease progression(*19*, *20*, *48*, *49*). The major challenge to the emerging field is the lack of ancestral transcriptional states in a reconstructed cell phylogeny, which prevents true longitudinal tracing of transcriptional dynamics(*50*). In this study, we resolved this problem and conducted, for the first time in a non-nematode organism, lineage-informed longitudinal analysis of cell state dynamics throughout development. Specifically, we reconstructed high-resolution zygote-to-larva developmental cell phylogenies in zebrafish. We then developed and validated LEAP, a computational framework for imputing ancestral transcriptional states of a reconstructed cell phylogeny. Subsequent phylogeny-informed longitudinal analyses revealed, in addition to gastrulation, a previously unappreciated wave of fate specializations during the hatching-to-larval transition. We also uncovered abundant incipient cell states with already-determined cell fates but minimal transcriptomic divergence, suggesting hidden developmental heterogeneity before overt transcriptional separation. Such heterogeneity may result from intrinsic molecular divergences (e.g., epigenetic changes) and/or extrinsic developmental context (e.g., spatial position), calling for additional modes of single-cell profiling to delineate this issue. These findings highlight cell phylogeny as a complementary framework to state-based atlases for resolving the historical and functional dimensions of cellular diversification(*23*). In conclusion, by resolving the ancestral states of a reconstructed cell phylogeny, this work paves the way for constructing lineage-informed cell atlases to fully understand cell state dynamics in complex organisms.

Despite these insights, several limitations or technical issues are worth discussing. First, the current scale of individual phylogenies limits the statistical power to resolve the full spectrum of phylogenetic states; expanding cell numbers per individual will be essential to enhance ancestral representation. Second, although the incipient phylogenetic states identified here show distinct terminal fate biases, our current data do not distinguish whether these biases arise from intrinsic molecular programs, spatial patterning, local environmental cues, or a combination of these factors. Because spatial information was not measured in this study, future integration with spatial transcriptomics, live imaging, or spatially resolved perturbation approaches will be required to map ancestral cells to their physical niches and to separate intrinsic factors from extrinsic context(*51–53*). Third, integrating scSMALT with modalities like scATAC-seq or single-cell methylation is needed to determine if phylogenetic states are coupled with epigenetic priming(*54*, *55*). Finally, enhancing ancestral cells recording at earlier stages, like gastrulation, remains a challenge due to barcode expression kinetics and total cell number in zebrafish, calling for future optimizations of the barcoding system^54,55^.

## Methods

### Construction of the 1-kb DNA barcode plasmids

The 1-kb DNA barcodes, HMF1 and HMF2, were designed based on previous mutation-readout studies in Drosophila and Saccharomyces cerevisiae(*16*). Both 1-kb barcodes contain 16 I-SceI binding motifs flanked by AID-preferred sequence contexts. Compared to the original 3-kb barcode used in Drosophila, this optimized shortened version features an 80-bp spacer between the 5′ ends of opposing I-SceI motifs and a 4-bp interval between their 3′ ends. The barcode cassettes were commercially synthesized (Azenta Life Sciences) with BamHI and NotI restriction sites incorporated at both ends. Subsequently, each barcode cassette was cloned into the 3′ untranslated region (UTR) downstream of a GFP coding sequence in a Tol2 vector(*56*, *57*). A T7 promoter was engineered between GFP and each barcode to facilitate downstream barcode recovery(*58*, *59*). These components were assembled to generate the dual-barcode transgenic construct Tol2-ubi:GFP-HMF2-pA-ubi:GFP-HMF1-pA, which will be available upon request. The complete sequences of HMF1 and HMF2 are provided in Supplementary Table 1.

### Generation of transgenic zebrafish lines

Wild-type AB zebrafish were maintained at 28.5 °C under a 14 h/10 h light/dark cycle. Tol2 transposase mRNA was synthesized in vitro using the HiScribe T7 High Yield RNA Synthesis Kit (New England Biolabs, E2060S) and purified using the Monarch RNA Cleanup Kit (New England Biolabs, T2040S). A mixture containing Tol2 transposase mRNA and the Tol2-ubi:GFP-HMF2-pA-ubi:GFP-HMF1-pA plasmid (at a final concentration of 12.5 ng/μL each) was microinjected into the yolk of one-cell stage AB wild-type embryos (approximately 2 nL per embryo). F0 embryos exhibiting ubiquitous GFP fluorescence were raised to adulthood. The transgenic status of potential founders was confirmed by outcrossing with wild-type AB fish; F1 founders producing 50% GFP-positive progeny—indicative of a single-copy insertion—were subsequently selected. F2 progeny from these single-copy founders were raised as stable transgenic lines and incrossed to establish a homozygous transgenic line. All animal procedures were approved by the Animal Research and Ethics Committee of Sun Yat-Sen University and conducted in accordance with the National Guidelines for the Care and Use of Laboratory Animals.

### Induction and validation of barcoding mutations

The AID-I-SceI-UGI fusion fragment of the SMALT system was PCR-amplified and cloned into a pCS2 vector downstream of a T7 promoter. Capped AID-I-SceI-UGI mRNA was transcribed in vitro using CleanCap® Reagent AG (TriLink BioTechnologies, N-7113-5) and N1-methylpseudouridine (TriLink BioTechnologies, N-1081-5) along with the HiScribe T7 kit (New England Biolabs, E2040S). Homozygous Tg(ubi:GFP-HMF2-pA-ubi:GFP-HMF1-pA) fish were outcrossed to wild-type AB fish, and the resulting one-cell stage embryos were microinjected with 300 pg of AID-I-SceI-UGI mRNA. Transgenic embryos displaying ubiquitous GFP expression and morphologically normal development were raised to specific developmental stages for analysis.

To assess the mutation accumulation rate over time, genomic DNA was extracted from pooled transgenic embryos at defined developmental stages: 2, 3, 4.7, 10, 24, and 30 hours post-fertilization (hpf) (n = 20 embryos per pool for stages before 10 hpf; n = 10 for later stages). Genomic DNA from 10 randomly selected, uninjected transgenic embryos at 24 hpf served as the negative control. The HMF1 and HMF2 barcodes were separately PCR-amplified and subcloned using a blunt-end cloning kit (CloneSmarter, C5851). Mutations were identified and quantified via Sanger sequencing of randomly selected individual clones. Cell numbers for the corresponding developmental stages were obtained from previous literature (*47*, *60*) where available, and supplemented by direct cell counting at 30 hpf in this study. For each stage, the approximate number of cell divisions was estimated as log2(N), where N denotes the total cell number. The resulting cell numbers and estimated division numbers are summarized in Supplementary Table 3.

### Preparation of single-cell suspensions for 7 dpf scSMALT zebrafish

Morphologically normal scSMALT zebrafish larvae at 7 dpf were individually transferred to microcentrifuge tubes, and the E3 medium was thoroughly removed. Immediately, 100 µL of 1× HBSS (Thermo Fisher Scientific, 14185052) containing 1× TrypLE Select (diluted from a 10× stock; Thermo Fisher Scientific, A1217701) was added. The tissue was mechanically dissociated via gentle pipetting, followed by a 30-second incubation at room temperature. This mechanical disruption process was repeated until no macroscopic tissue fragments remained. Dissociation efficacy was verified by examining a 1 µL aliquot under a microscope to confirm the single-cell proportion and structural integrity. To stop the dissociation reaction, 900 µL of ice-cold 1× HBSS supplemented with 0.05% BSA was added, and the cells were gently resuspended. The cell suspension was passed through a 35-µm cell strainer pre-wetted with 500 µL of 1× HBSS containing 0.05% BSA. An additional 200 µL of the same buffer was used to rinse the strainer. The filtered single-cell suspension was centrifuged at 500 × g for 5 minutes at 4 °C. The supernatant was carefully aspirated to avoid disturbing the cell pellet, leaving approximately 50 µL of suspension. Cell concentration was determined using a hemocytometer, and the suspension was adjusted to a final concentration of ∼1,000 cells/µL. Approximately 15,000 cells per sample were loaded onto the Chromium Next GEM Single Cell 3ʹ v3.1 platform (10x Genomics) according to the manufacturer’s protocol. Gene expression libraries were constructed and subsequently sequenced on Illumina HiSeq or NovaSeq platforms.

### Analysis for 10x scRNA-seq data

Alignment of sequencing reads and processing into a digital gene expression matrix were performed using Cell Ranger (v.6.1.2), including the aligner STAR (v.2.7.2a)(*61*), with standard parameters. To specifically quantify mature mRNA transcripts, the *--include-introns* parameter was set to false. The data were aligned against the zebrafish reference genome GRCz11 (Ensembl release 105). The *--expect-cells* parameter was set to 5,000–20,000 based on the number of cells loaded per sample. Gene expression matrices were processed and analyzed using the Seurat R package (v5)(*62*, *63*). For initial quality control, cells were filtered based on the number of detected genes (nFeature_RNA > 200) and the percentage of mitochondrial reads (percent.mt < 20%) to ensure high-quality transcriptomes. Potentially confounding doublets were identified and excluded using DoubletFinder(*64*), and only predicted singlets were retained for downstream analysis. Following doublet removal, the data were normalized and variance-stabilized using *SCTransform*(*65*). To mitigate batch effects across multiple samples, we utilized the Harmony algorithm for data integration. The integrated dataset was then subjected to principal component analysis (PCA), followed by non-linear dimensionality reduction via UMAP embedding(*66*). Cluster-specific marker genes were identified using the *FindMarkers* function (Wilcoxon rank-sum test). Cell-type identities were manually annotated according to the expression of these top-ranking markers, cross-referenced with established literature and the ZFIN database (*30–33*). After cell-type annotation, the top 50 marker genes were identified for each annotated cell type using the same marker detection framework. The union of these cell-type markers, after removing redundant genes, was used as the cell-type marker set for comparison with LUGs in Fig. 3e.

### Targeted amplification of 1-kb barcodes from 10x full-length cDNA libraries

To integrate cellular identity with lineage history within the scSMALT framework, we developed a high-specificity enrichment strategy to recover the 1-kb lineage barcodes. In the transgenic lines, these barcodes are expressed as the 3′ UTR of GFP under the control of the ubiquitous ubi promoter. During scRNA-seq library preparation, the polyadenylated barcodes are captured by 10x Genomics gel beads, inextricably linking cell-type transcriptomes to lineage-specific mutational profiles via shared 10x cell barcodes. To selectively enrich these barcodes from the full-length cDNA pool, an in vitro transcription (IVT)-mediated approach was employed. Specifically, 8 µL of the 10x full-length cDNA was used as the template for IVT with the HiScribe T7 High Yield RNA Synthesis Kit at 37 °C for 2 hours. Following transcription, the residual cDNA template was degraded using DNase I (New England Biolabs), yielding the barcode sequences exclusively in RNA form. The in vitro-transcribed RNA was purified using the Monarch RNA Cleanup Kit. First-strand cDNA was subsequently synthesized using the RevertAid cDNA Synthesis Kit (Thermo Fisher Scientific, K1622) with a Read 1-specific primer to target 10x-barcoded molecules selectively.

The final 1-kb amplicons were generated via PCR using KOD One DNA Polymerase (Toyobo) with primers targeting the partial Read 1 sequence and the respective HMF1/HMF2 forward sequences. To minimize PCR-induced chimeras and maintain high molecular fidelity, the number of amplification cycles was empirically optimized (typically 18–25 cycles) to yield approximately 1 µg of target DNA required for PacBio SMRT sequencing. The PCR products were purified utilizing a 0.6× ratio of SPRIselect beads (Beckman Coulter) to remove non-specific short fragments. All primer sequences are detailed in Supplementary Table 2. The integrity of the amplicons was verified via blunt-end cloning and Sanger sequencing before PacBio SMRT sequencing.

### SMRT sequencing and data processing

One microgram of the PCR 1-kb barcode amplicons was utilized for SMRTbell library preparation (PacBio Sequel II), following the standard SMRT sequencing protocol. Sequencing runs were conducted using one SMRT cell for each SMRTbell library. For the PacBio Sequel II subread group, circular consensus sequences (CCS) were generated by pbccs v.6.4.0 with the parameters *--min-length=1000 --min-passes=3*.

Following CCS generation, index sequences, including 10x cell barcodes and unique molecular identifiers (UMIs), were extracted using the SeqKit toolkit(*67*). To mitigate sequencing noise, index sequences were clustered with usearch, and dominant groups with a relative frequency over 50% were selected for further analysis. A two-stage consensus merging process was then applied to the index-specific HMF sequences using usearch: the first stage merged sequences based on entire index sequences, while the second stage merged the output sequences based on cell barcodes alone. Sequences shorter than 600 bp were considered low-quality and excluded from subsequent analysis.

Finally, the remaining HMF sequences were aligned to the reference genome using minimap2(*68*) with the *-ax map-pb* parameter, and mutations were identified using bcftools *mpileup*(*69*). To ensure high confidence in the results, variants were filtered by a quality score (QUAL) exceeding 20, with the additional requirement that the frequency of the dominant alternative allele must be greater than 50%.

### Cell Phylogenies reconstruction

Maximum-likelihood cell phylogenies were reconstructed using IQ-TREE. To ensure an optimal evolutionary model, the best-fit substitution model for each dataset was determined with the selection criteria based on the Bayesian Information Criterion (BIC). Subsequently, ML trees were inferred using the selected model, with the wildtype HMF sequence designated as the outgroup (*-o Ref*). Parameters for tree inference included ancestral state reconstruction (*-asr*), 1,000 SH-like approximate likelihood ratio test iterations (*-alrt 1000*), and 1,000 ultrafast bootstrap replicates (*-bb 1000*) with a maximum of 1,000 iterations (*-nm 1000*). The cell type annotations were integrated by mapping the 16-nt 10x Genomics cell barcodes to the tree tips.

To refine the topology and eliminate uncertainty associated with low-confidence nodes, branches with near-zero length were collapsed into polytomies using the TarCA package(*25*). For a fully resolved phylogeny, the number of internal nodes should be equal to the number of terminal cells minus 1. Then the degree of phylogenetic resolution was quantified as the proportion of resolved ancestral division events, calculated as *N*/(*T*-1), where *N* represents the number of internal nodes and *T* represents the number of terminal cells.

### Transcription and lineage kinship analysis

Lineage-associated phylogenetic similarity within cell types was quantified using the coalescent rate (CR) calculated via the *Np_Estimator* function of the TarCA package. A higher CR represents a closer phylogenetic similarity, while the lower CR is opposite. Analysis was restricted to cell types represented by at least 20 cells for an individual phylogeny. The obtained CR values were compared against a null distribution generated from 1,000 random permutations.

Transcriptomic similarity was assessed using Euclidean distances in the Harmony embedding space. For a focal cell *i*, transcriptional divergence (*D_i_*) was measured as the average distance between cell *i* and its identified phylogenetic relatives (e.g. sisters). A background divergence value (*R_i_*) was obtained through random label shuffling of the relative population. The transcriptional similarity score was calculated as 1-*D_i_*/*R_i_*. This metric was then applied iteratively to evaluate relationships across increasing phylogenetic depths, extending from immediate sister cells to first- through third-order cousins. Finally, the lineage coupling index for a given cell type was defined as the sum of the average transcription similarity across the four phylogenetic relatives.

### Identification of LUGs

The identification of lineage-committed upregulated Genes (LUGs) followed the methodology previously described(*25*). Briefly, transcriptomic profiles were first imputed using SAVER(*70*). Genes expressed in fewer than 10% of total cells were excluded to minimize noise from low-abundance transcripts. For the remaining genes, cells with expression levels exceeding the 90th percentile were classified as an upregulated population (assigned a value of 1), while all other cells were assigned 0.

Lineage association for each gene was then quantified via the coalescent rate calculated with the *LEU_Estimator* function of the TarCA package. Statistical significance was established by comparing observed CR values against a null distribution generated from 1,000 random permutations. Genes with fewer than 30 representative cells per phylogeny or an empirical P-value exceeding 0.01 were excluded. The empirical P-values were further aggregated across all phylogenies using the harmonic average. Genes with a resulting harmonic P-value < 0.01 were classified as highly confident LUGs.

Overlap analysis between LUGs and markers was conducted across two layers. Specifically, the top 50 cell-type-specific markers were used for this comparison.

Additionally, germ layer marker analysis was restricted to the surface ectoderm, neuroectoderm, mesoderm, and endoderm.

### LUG-based binarization of transcriptomic matrices

For each highly confident LUG, the activation nodes were located with the most recent common ancestor (MRCA) of the cells exhibiting an enriched upregulation signal, as determined by the TarCA package. The activation node was defined as the root of the subclade where the upregulation event was initiated. A binary vector was generated for each LUG by assigning a value of 1 to all descendants, including all relevant internal nodes and terminal nodes, of the respective MRCA and 0 to the rest. These individual vectors were subsequently integrated to construct a binary matrix representing the lineage-specific activation landscape.

### LEAP computational framework

The LEAP computational framework utilized two matrices: a query matrix and a reference matrix. The query matrix was generated by the LUG-based pipeline, where each row represented a LUG and each column represented an internal node. The reference matrix is derived from single-cell transcriptomic data with the same columns and rows as the query matrix. Specifically, cells with expression levels exceeding the 90th percentile were assigned a value of 1, while the remainder were assigned 0. These matrices were then processed using the Seurat pipeline, including *NormalizeData*, *ScaleData*, *RunPCA*, and *RunUMAP*. Finally, labels from the reference data were projected onto the query data using the *TransferData* function. Specifically, the transferred labels were customized according to the reference dataset.

### Integration of query and reference data

The query and reference objects were first joined using the merge function, followed by RunUMAP to visualize the joint manifold. This resultant object was termed the "DirectJoin" object. Alternatively, the query and reference objects were integrated using either *CCAIntegration* or *HarmonyIntegration* to generate the "Integration" object. The consistency between the query and reference datasets was quantified using the inverse Local Inverse Simpson’s Index (iLISI). An iLISI score approaching the theoretical maximum indicates well-mixed populations. Note that the maximum iLISI is data-dependent; for two datasets, the theoretical maximum is 2, but the observed maximum can be slightly lower depending on the number of cells in each batch. To provide a rigorous baseline, we also assessed the empirical maximum by performing in silico mixing of the two datasets.

### LEAP validation on *C. elegans*

The reference phylogenetic tree was derived from the Sulston et al. *C. elegans* embryogenesis dataset(*71*), and the lineage-resolved single-cell RNA sequencing atlas for *C. elegans* was obtained from publicly available datasets(*36*). LUG identification was performed using the TarCA package, and the states of internal nodes were inferred following the LEAP framework.

The consistency between the inferred internal nodes and the ground truth was assessed from three perspectives. First, we focused on the five primary lineages, including AB, C, D, E, and MS, within the phylogeny and constructed a confusion matrix to compare the inferred states against the ground truth. Second, we evaluated the accuracy of individual cell lineage assignments. Given that the lineage identities in the reference atlas were degenerate, an inferred state was considered consistent if it represented a perfect match (Dist=0) or allowed for a single mismatch (Dist=1) against the corresponding ancestral cells, with background levels established via 1,000 permutations. Third, we validated the consistency of the inferred states across four broad developmental cell types, including epidermis (Epi), neurons (Neu), muscle (Mus), and intestine (Int).

### LEAP application on zebrafish

We applied the TarCA package to 30 zebrafish phylogenetic trees and their corresponding single-cell RNA sequencing datasets to extract LUGs and to reconstruct transcriptomic profiles of internal nodes. For each resolved internal node, we calculated its distance from the root, the number of descendants, and the diversity of descendants (defined as the number of distinct terminal cell types within the clade). Subsequently, internal node state inference was performed using the LEAP framework. Two features were transferred from the reference atlas: the developmental stage (denoted by the stage.group column) and cell type identity (denoted by the identity.super column). For each cell type, the inferred transcriptome was represented by the average expression levels of LUGs at the internal nodes, while the transcriptome of actual cells was represented by the average expression levels of cells derived from the reference dataset.

### Identification of cell-type-specific LUGs

For each cell type with at least 30 cases, we quantified the specificity of each LUG by calculating a marker score, defined as *P_t_*/(1-*P_b_*), where *P_t_* denotes the proportion of the activation events within the focal cell type, while *P_b_* represents the proportion of the activation events across all other cell types. Finally, candidate LUGs for each cell type were ranked in descending order of their enrichment scores, enabling the selection of high-confidence markers for subsequent functional validation.

### CRISPR–Cas9 mutagenesis in zebrafish embryos

The top five ranking LUGs for ‘neurons’ were selected for candidate knockouts. For CRISPR-Cas9-mediated mutagenesis, gRNAs were identified via CHOPCHOP(*72*) , and the crRNA sequences used for each target gene are listed in Supplementary Table **a.** 5. We prepared ribonucleoprotein (RNP) complexes following a high-efficiency zebrafish protocol described previously(*45*). Briefly, Alt-R crRNA and tracrRNA (IDT) were hybridized at a 1:1 molar ratio to form a 50 µM duplex. Recombinant Cas9 protein (V3, IDT) was prepared at 25 µM in a specialized injection buffer (20 mM HEPES-NaOH, 350 mM KCl, and 20% glycerol). Before injection, RNPs were assembled by mixing the crRNA:tracrRNA duplex (equimolar pools for multiplexing) with Cas9 protein at a 1:1 volume ratio to ensure optimal cleavage activity. The RNP mixture was incubated at 37 °C for 5 minutes to facilitate complex formation and subsequently maintained at room temperature. Approximately 2 nl of the RNP solution was microinjected into the cytoplasm of wild-type zebrafish embryos at the one-cell stage.

### Genotyping of F0 Crispants

To evaluate gene-editing efficiency, genomic DNA (gDNA) was extracted from F0 embryos via alkaline lysis following established protocols(*45*, *46*, *73*). Briefly, at 48 hours post-fertilization (hpf), pools of five randomly selected F0 embryos per sgRNA treatment were lysed in 100 μL of 50 mM NaOH and incubated at 95°C for 10 min. The lysates were neutralized with 10 μL of 1 M Tris-HCl (pH 8.0), and 2 μL of the resulting crude gDNA solution was used as a template for PCR. Target regions flanking the sgRNA sites were amplified using specific primers designed via CHOPCHOP (v3)(*72*). Genomic DNA from 48-hpf wild-type (WT) embryos served as the control. PCR amplicons were purified through gel extraction and subjected to Sanger sequencing. For characterization of indel variants, the purified PCR products were cloned using a blunt-end cloning kit (CloneSmarter, #C5851) according to the manufacturer’s instructions. 20 clones per sample were randomly selected for Sanger sequencing to validate the mutation efficiency.

### Preparation of single-cell suspensions for 2dpf F0 crispants

F0 crispant embryos targeting LUGs were generated via CRISPR–Cas9 mutagenesis of embryos derived from crosses between the AB and Tu zebrafish lines. To maximize throughput and SNP divergence, pools of 4 to 6 embryos were assembled, with each individual sourced from a distinct parental pair. At 2 dpf, the pooled embryos were dechorionated using a 0.5 mg/mL pronase solution (5 mL per dish), following a previously described protocol(*74*). To remove the yolk, the ‘dechorionated embryos were transferred into 0.5 mL of ice-cold deyolking buffer (55 mM NaCl, 1.8 mM KCl, 1.25 mM NaHCO3). The yolk sacs were mechanically ruptured via gentle pipetting for 30 seconds, and the samples were subsequently centrifuged at 500 × g for 1 minute at 4 °C. The deyolked embryonic tissue was transferred to a microcentrifuge tube, and the residual buffer was removed as completely as possible. Subsequent tissue dissociation and single-cell suspension purification were performed as described above for the 7 dpf larvae, albeit with a reduced enzymatic lysis time when no visible large tissue clusters remained. Single-cell RNA-seq libraries were then constructed utilizing the DNBelab C4 platform in strict adherence to the manufacturer’s guidelines.

## Genetic demultiplexing of LUG KO scRNA-seq data

To achieve high-throughput profiling while minimizing batch effects, multiple individuals—including wild-type (WT) and various LUG crispants—were pooled prior to single-cell library preparation using the DNBelab C Series platform. Following sequencing, raw reads were processed and aligned to the zebrafish reference genome (GRCz11) using dnbc4tools (v2.1.3) to generate cell-associated BAM files (https://github.com/MGI-tech-bioinformatics/DNBelab_C_Series_HT_scRNA-analysis-software). To ensure interoperability with the Souporcell pipeline(*75*), we employed a custom stream-processing script to adjust the BAM metadata. Specifically, we performed a tag-substitution where the DNB-specific droplet barcode tag (DB) was converted to the standard cell barcode tag (CB). Genetic demultiplexing was subsequently performed by executing the souporcell_pipeline.py with the cluster parameter (k) set to the exact number of individuals pooled in each library. The pipeline integrated FreeBayes for de novo variant calling and Vartrix for allele quantification per cell. For each cell, Souporcell calculated the log-likelihood of its membership within each genotype cluster, enabling the classification of cells into singletons, cross-genotype doublets, or unassigned cells. To ensure data integrity, only cells identified as singletons with high-confidence genotype assignments were retained for downstream analysis. Doublets and cells with ambiguous assignments were excluded from the final dataset.

### scRNA-seq analysis of 2dpf LUG KO scRNA-seq data

For the 2dpf LUG KO dataset, single-cell transcriptomes from wild-type (WT) and five specific LUG crispants (*dnajc5aa, smarcd1, fam168b, pdcl, and tra2b*) were integrated for comparative analysis. For initial quality control, cells were filtered based on the number of detected genes (nFeature_RNA > 200) and the percentage of mitochondrial reads (percent.mt < 20%) to ensure high-quality transcriptomes. To ensure the accuracy of individual assignments and minimize contamination, the dataset was subsetted to include only high-confidence singletons identified by the Souporcell genetic demultiplexing pipeline. The data were then normalized and variance-stabilized using *SCTransform*. To achieve precise and consistent cell-type annotation with the 7dpf scSMALT dataset, we performed reference-based label transfer. The 2dpf query dataset was mapped onto the 2dpf zebrafish developmental reference atlas (23 daniocell)(*30*) using the *FindTransferAnchors* and *TransferData* functions in Seurat (v5)(*63*), utilizing the top 50 dimensions. Predicted identities were then integrated into the metadata. For visualization and unsupervised clustering, we performed Principal Component Analysis (PCA) on the integrated data. Non-linear dimensionality reduction was executed via UMAP embedding based on the first 50 principal components. Cell clustering was performed using the *FindNeighbors* and *FindClusters* functions to capture fine-grained cellular heterogeneity across the wild-type and mutant individuals. Daniocell single-cell transcriptomic reference datasets used in this study are publicly available at https://daniocell.nichd.nih.gov/.

### Statistical assessment of cell-abundance changes

To mitigate technical artifacts arising from variations in sequencing depth and total cell counts, we implemented a geometric mean-based normalization approach following a previously published algorithm(*46*). Specifically, we calculated the geometric mean of total cell counts across all embryos and defined the size factor for each sample as the ratio of its total cell count to this global geometric mean. To ensure normalization robustness, we prioritized cell types with high compositional stability, defined by the lowest coefficient of variation across replicates, to refine the size-factor estimation. Raw cell counts were then normalized by these size factors to obtain the normalized cell number.

For each cell type with an overall cell number over 500, we modeled the normalized counts using a beta-binomial generalized linear model, with the size factors included as an offset term. The model is defined by the following equations:

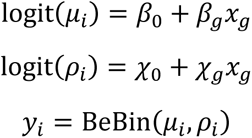

In this framework, *y_i_* represents the normalized counts of cell type *i*, modeled with a mean *μ_i_* and a "litter effect" *ρ_i_*. We modeled both parameters as functions of the genotype *x_g_* to account for potential differences in variability between wild-type and crispant embryos. The coefficients *β_g_* and *χ_g_* quantify the effect of the knockout on the mean abundance and the overdispersion, respectively; *β*_0_ and *χ*_0_ are the corresponding intercepts. We also included the number of periderm cells as a nuisance term to control for overall animal size. Models were implemented using the VGAM package(*76*). The effect size of the knockout was quantified as the Odds Ratio (OR, i.e. exp(*β_g_*)). Statistical significance was assessed via the Wald test on the genotype coefficient *β_g_*, and *P*-values were adjusted for multiple testing using the Benjamini-Hochberg (BH) procedure to control the false discovery rate (FDR).

### Quantification of differentiation potency

Differentiation potency for each internal node within the lineage trees was quantified based on the composition of its descendant cell types. For a given internal node, let *q_i_* represent the relative proportion of its total descendants belonging to the *i*-th cell type. The differentiation potency of this internal node was defined by Shannon’s entropy as:

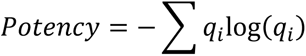

To identify lineage-specific differentiation tendencies, we performed a three-layer statistical validation on internal-terminal cell type pairs. Analysis was restricted to internal cell types represented by at least 20 internal nodes across the reconstructed trees. First, for each qualifying internal cell type, we extracted all corresponding terminal cells and calculated the observed proportion of each terminal cell type. The lineage association strength for a specific internal-terminal pair was defined as the average relative proportion of the focal terminal cell type across all qualifying internal nodes. We then implemented a permutation test with 1,000 randomized simulations, where the terminal cell type annotations were shuffled. A one-sample Wilcoxon signed-rank test was applied to compare the observed association strength against the null distribution, and the resulting *P*-values were adjusted for multiple testing using the BH procedure to control the FDR. Lineage pairs exhibiting an FDR ≥0.05 or a fold change ≤ 1.25 compared to the permutation background were filtered out. Second, we conducted a bootstrapping procedure with 1,000 independent replicates to assess the architectural robustness of the lineages. Pairs with a bootstrap support value ≤0.8 were discarded. Third, for the remaining cell types, we calculated the relative proportion of terminal cells within a specific cell type that were derived from the focal internal cells. Lineage pairs with a contributing proportion ≤20% were excluded from further analysis. The remaining pairs were defined as highly confident, statistically robust pairs of internal-terminal cell types.

### Identification of Phylogenetic and Transcriptional States

For the phylogenetic state analysis, a count matrix was constructed based on the terminal cell components. Within this matrix, each row corresponded to an individual internal node, and each column represented a high-confidence, statistically enriched terminal cell type. An entry of {0, 1, 2, …} denotes the absolute abundance of each terminal cell type derived from the focal internal node. Internal nodes associated with fewer than three terminal cells were excluded from downstream analysis. The matrix rows were then normalized into relative proportions. The normalized data were scaled, and dimensionality reduction was performed sequentially using PCA and UMAP. Finally, we applied shared nearest neighbor (SNN) density-based clustering using the dbscan(*77*) package to define distinct subpopulations of internal nodes, which are characterized by their specific phylogenetic states.

For the transcriptional state analysis, a binary matrix was constructed from the LUG-based binarization of the raw transcriptomic data. Each row represented an individual internal node, and each column corresponded to a specific LUG, where an entry of 1 indicated gene activation and 0 denoted the absence of activation. Internal nodes exhibiting fewer than three gene activation events were filtered out. The remaining binary matrix was converted into a Seurat object and processed through the standard pipeline, including data normalization, scaling, and dimensionality reduction was performed sequentially using PCA and UMAP to construct a transcriptional embedding space for the internal nodes.

### Assessment of Phylogenetic-State-Specific Differentiation Bias

For each identified subcluster in phylogenetic state, we compared the relative proportion of a focal terminal cell type within that subcluster against its counterpart across all remaining subclusters. Specifically, for each combination of subcluster and terminal cell type, a two-tailed Wilcoxon rank-sum test was performed to evaluate whether the internal nodes within the focal subcluster exhibited a significantly higher or lower differentiation propensity compared to the background nodes outside of it. Concurrently, the effect size of this differentiation bias was quantified using a log_2_-transformed fold change, calculated as the ratio of the average proportion within the cluster to the mean proportion in all other clusters.

## Data and code availability

Raw sequencing data have been deposited in the Genome Sequence Archive (GSA) under the National Genomics Data Center (NGDC), associated with BioProject accession PRJCA064592 (https://ngdc.cncb.ac.cn/). The code used for data processing and analysis is available at GitHub (https://github.com/shadowdeng1994/LEAP and https://github.com/shadowdeng1994/LEAP_sourcedata).

## Acknowledgements

This work was supported by the National Key R&D Program of China (2021YFA1302500 and 2021YFA1302501), the National Natural Science Foundation of China (32293190, 32293191, 12526210 and 32500524).

## Supplementary Materials

**Fig. S1.**
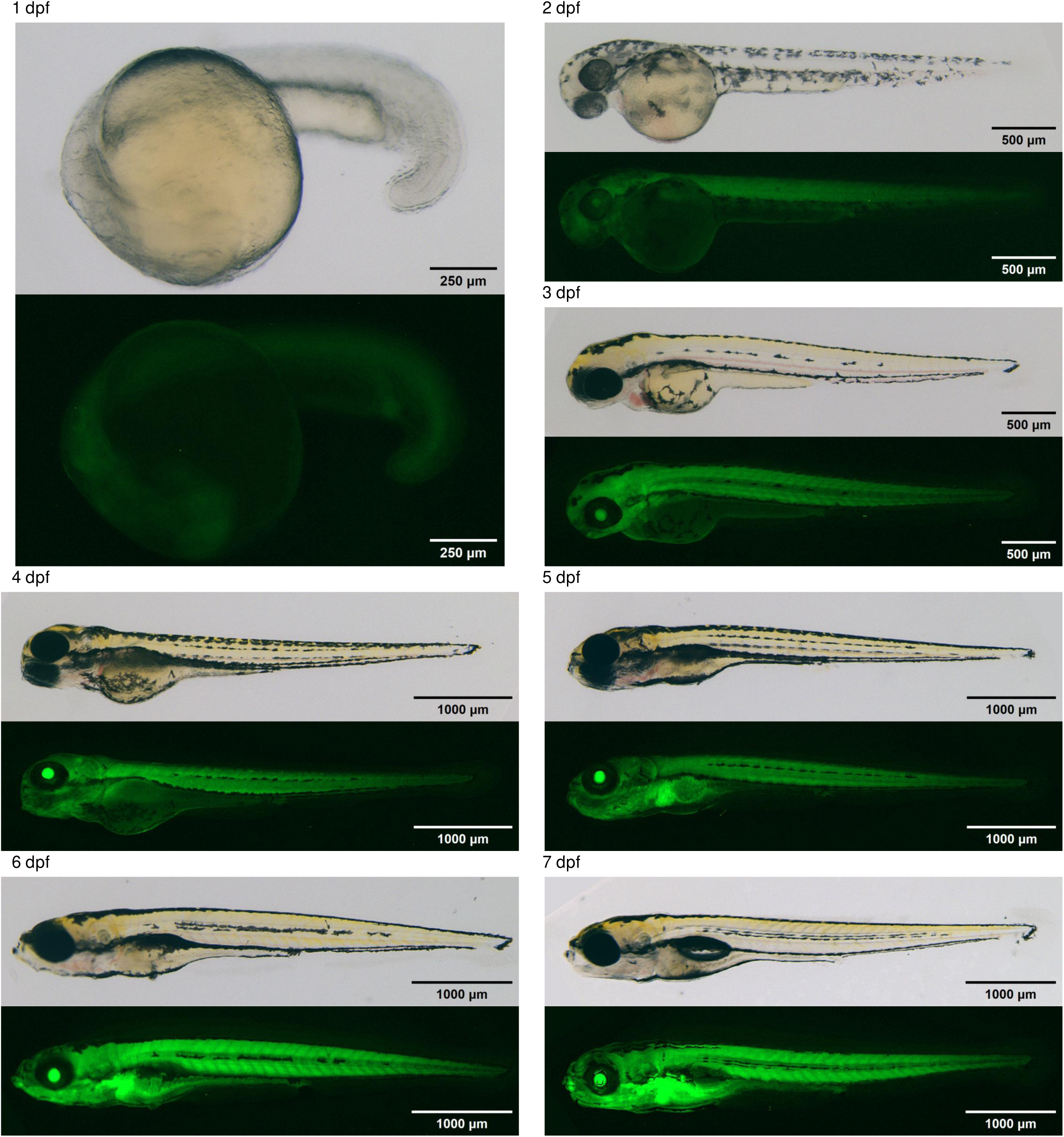
scSMALT transgenic line. Representative bright-field and fluorescence images of ScSMALT transgenic zebrafish larvae from 1 to 7 days post-fertilization (dpf). For each developmental stage, the upper image shows the bright-field view and the lower image shows the corresponding fluorescence signal. Scale bars: 250 μm for 1 dpf, 500 μm for 2–3 dpf, and 1000 μm for 4–7 dpf.

**Fig. S2.**
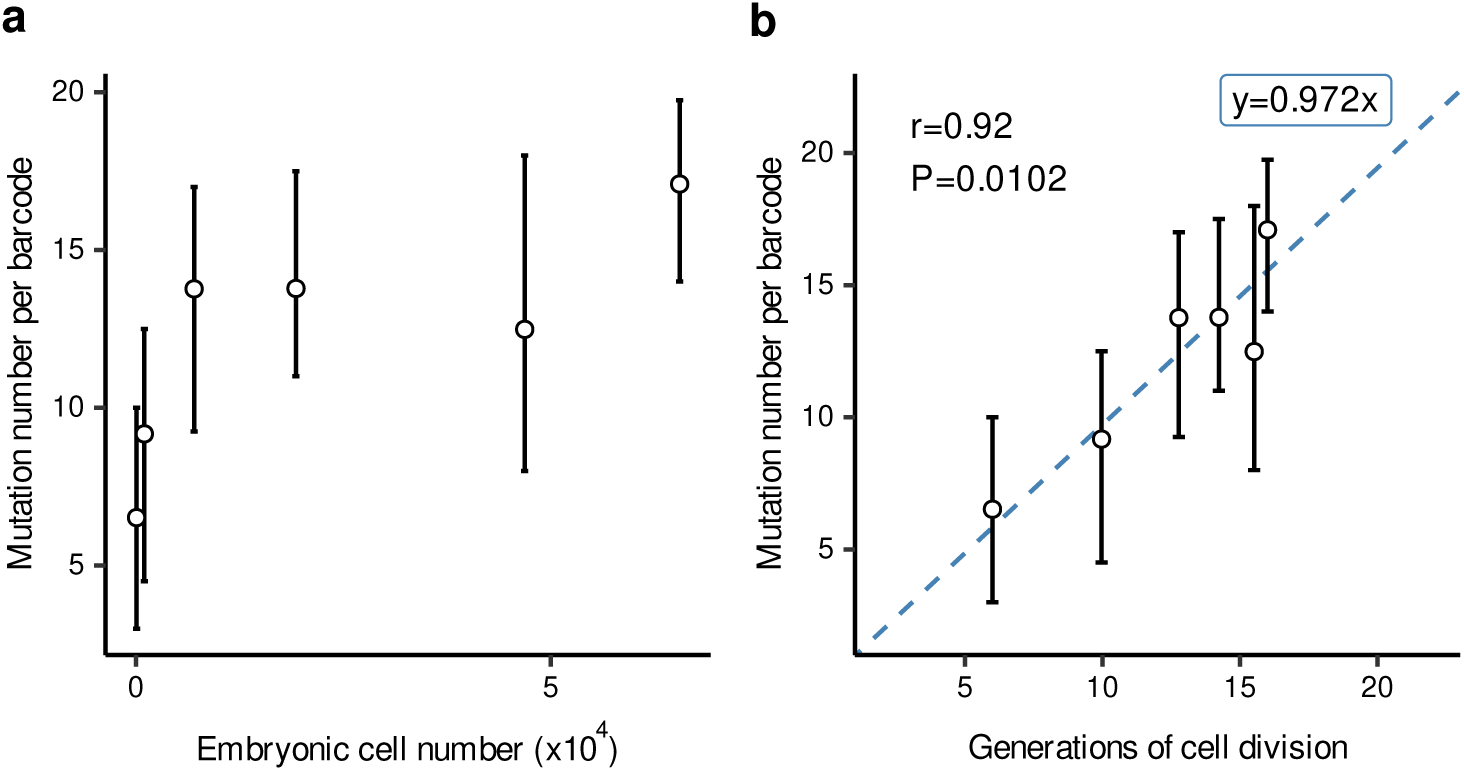
Relationship between mutation accumulation, population size, and total cell divisions. **a.** Distribution of mutation number per 1-kb barcode across developmental stages with corresponding estimated embryonic cell numbers. (as referenced in Fig. 1c). Boxplots show the number of mutations per barcode at each time point. Cell numbers for each stage are estimated based on the literature. **b.** Correlation between the estimated number of cell divisions and the number of mutations per barcode. The number of cell divisions is calculated as the log2 transformed of the total cell count at each developmental stage. The dashed line and Pearson correlation coefficient (r) illustrate the linear regression.

**Fig. S3.**
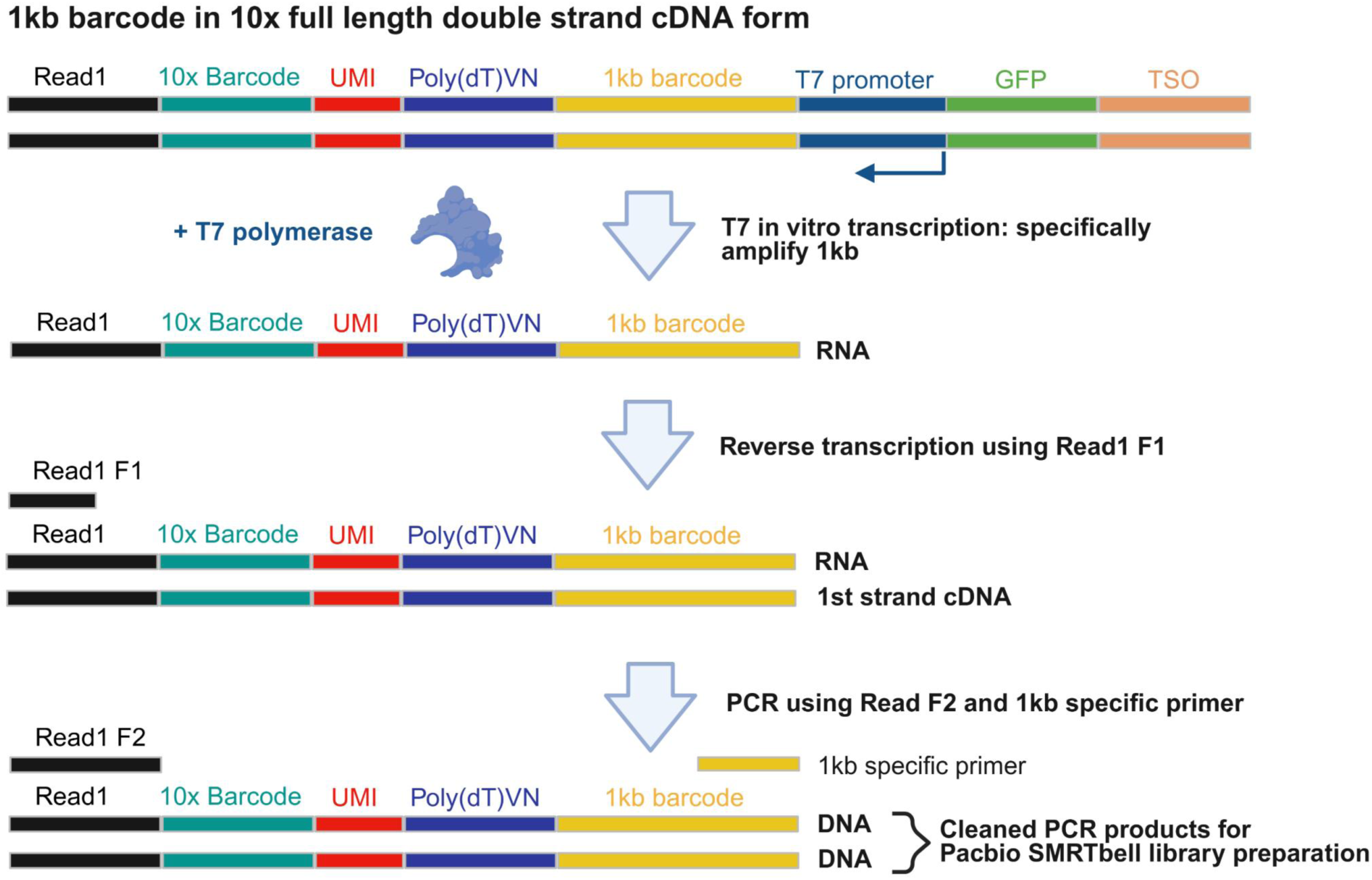
Workflow of T7-assisted target amplification for lineage barcodes. Schematic illustrating the step-by-step procedure to recover 1-kb barcodes from 10x Genomics full-length cDNA libraries selectively. The workflow includes: (1) in vitro transcription (IVT) of the T7-promoter-driven barcodes; (2) reverse transcription using a Read1-specific primer to target 10x-barcoded molecules; and (3) PCR amplification using high-fidelity DNA polymerase to generate final amplicons for PacBio SMRT sequencing.

**Fig. S4.**
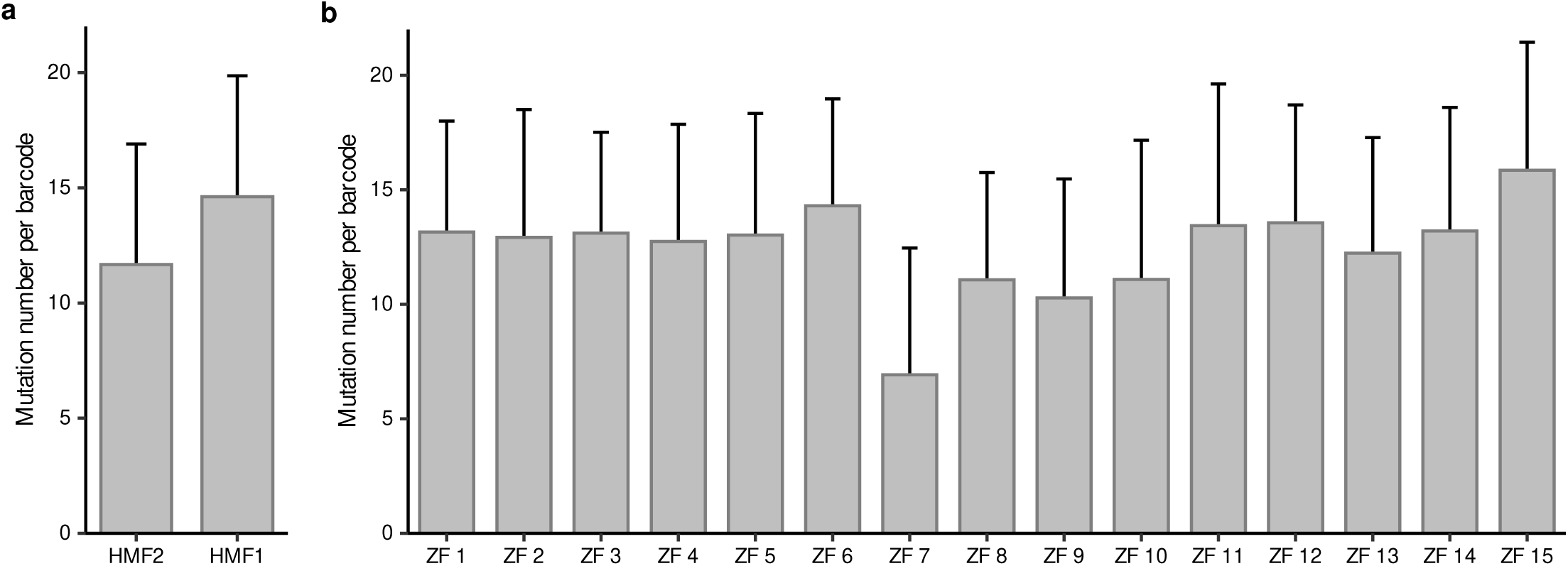
Mutation number per 1-kb barcode from 10x scRNA-seq cDNA. **a.** Distribution of mutation number per barcode for HMF1 and HMF2. Boxplots show the mutation number identified across all cells with recovered HMF1 or HMF2 1-kb barcode. **b.** Distribution of mutation number per barcode across 15 individual larvae. Boxplots show the mutation number for cells with recovered HMF1 or HMF2 1-kb barcode within each individual.

**Fig. S5.**
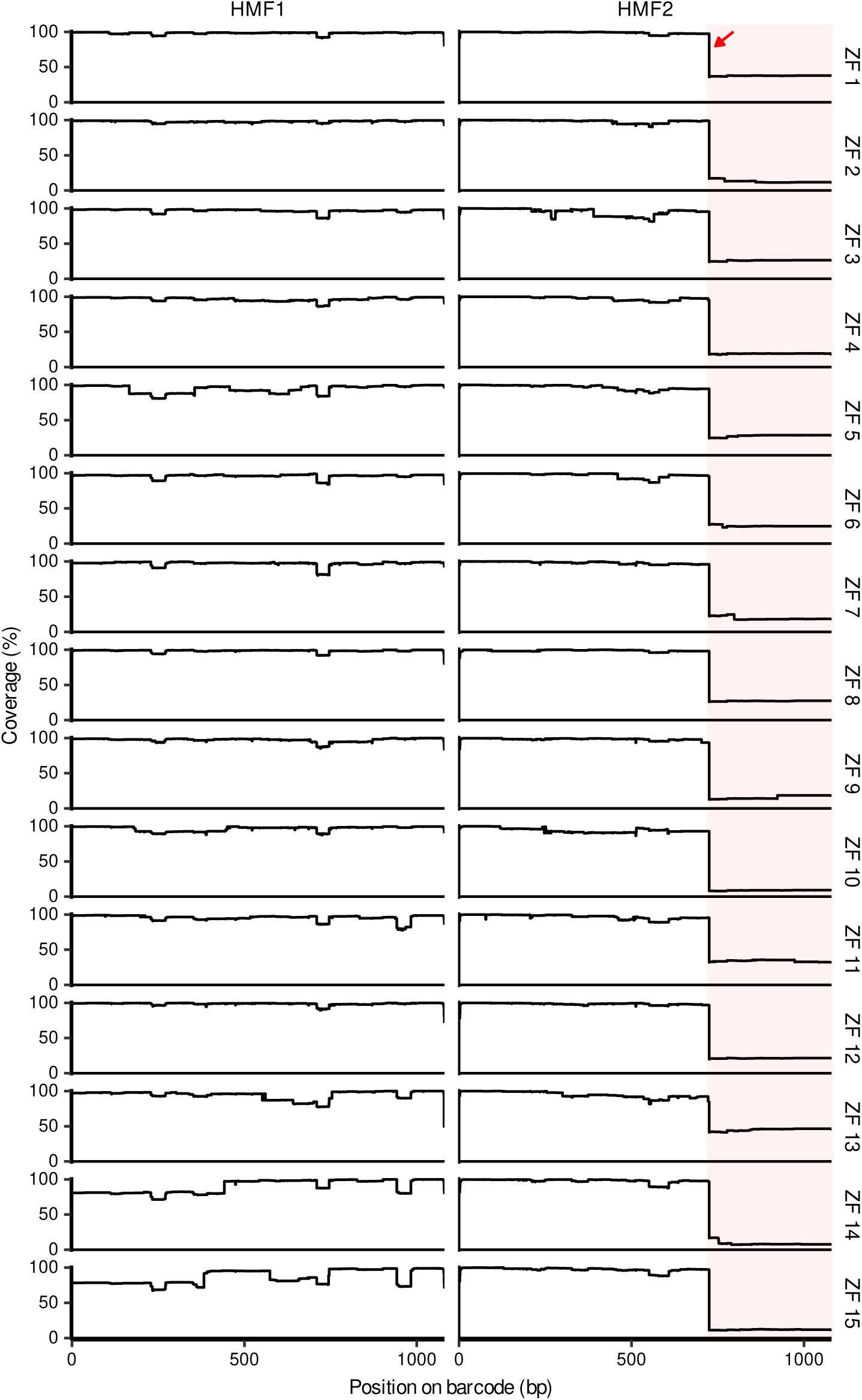
1-kb barcode coverage from 10x scRNA-seq cDNA library. Coverage profiles of amplified 1-kb barcode sequences from 10x scRNA-seq cDNA for 15 zebrafish individuals. For each individual, coverage across the barcode is shown separately for HMF1 and HMF2. The x-axis indicates the position on the barcode, and the y-axis indicates the percentage of cells with recovered HMF1 or HMF2 sequences at each barcode position. The shaded area marks the distal barcode region with reduced coverage in HMF2.

**Fig. S6.**
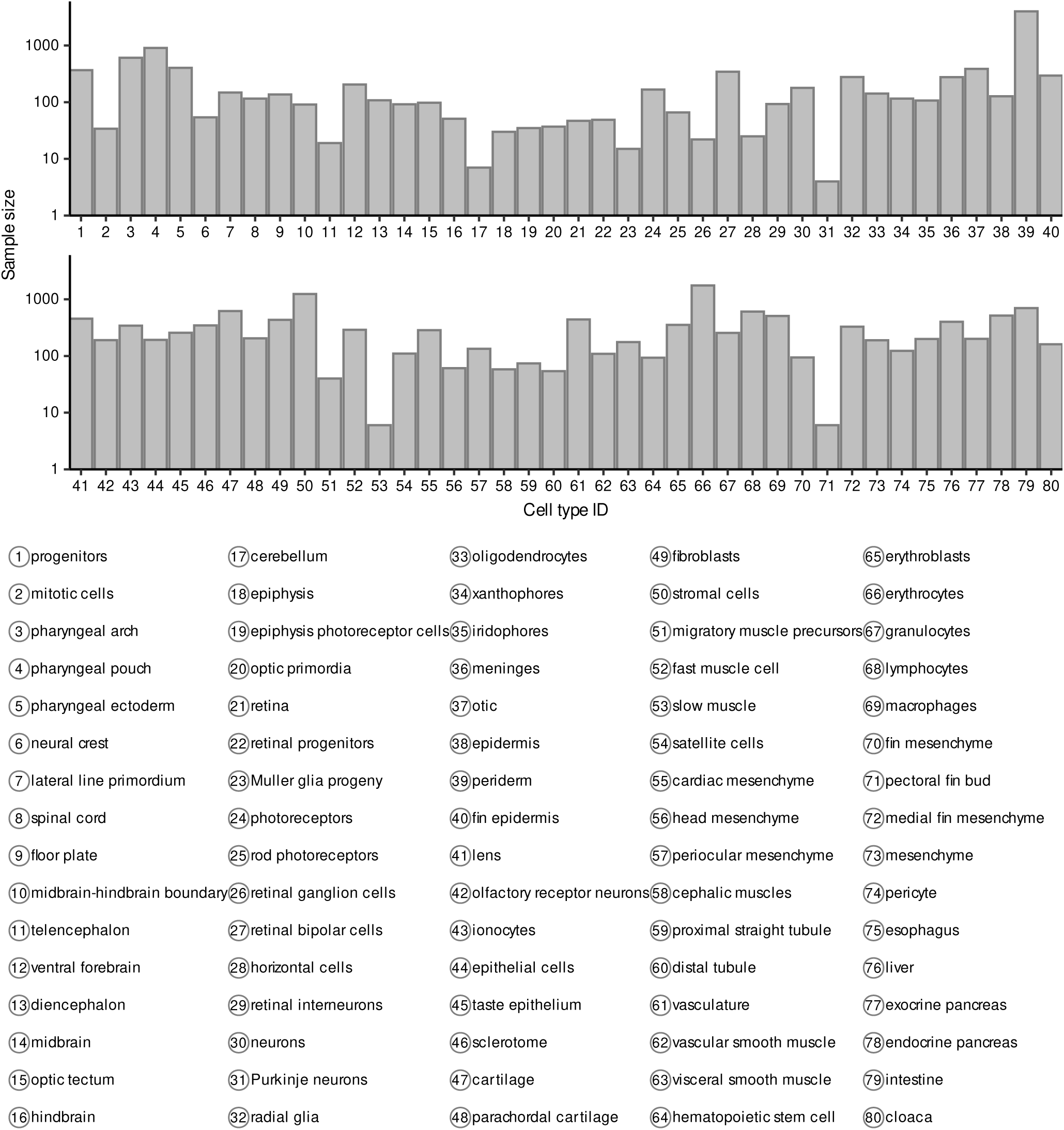
Number of cells recovered with a 1-kb barcode across 80 annotated cell types. Bar plot showing the number of cells recovered with a 1-kb barcode across the 80 annotated cell types. Each bar represents a distinct cell type, with the height corresponding to the total number of cells recovered with a 1-kb barcode. The y-axis is displayed on a logarithmic scale.

**Fig. S7.**
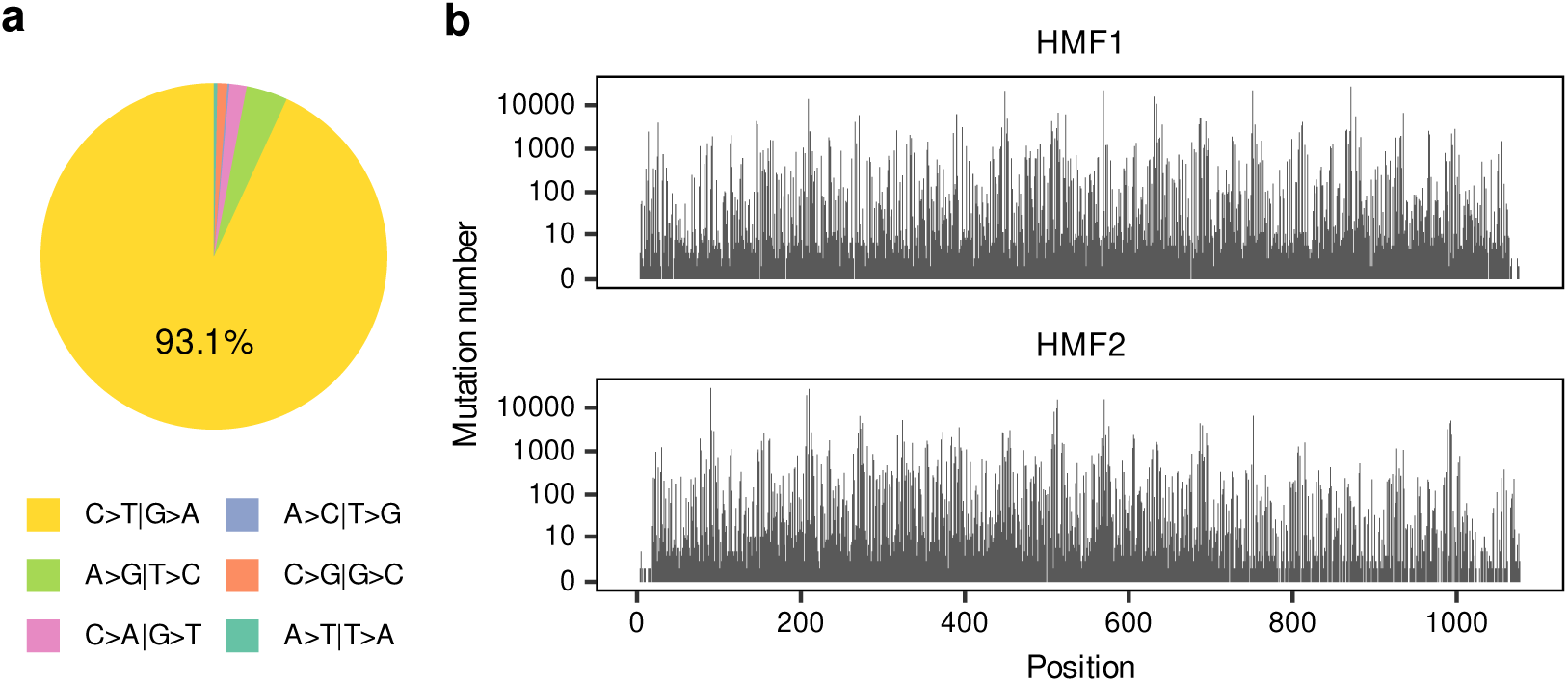
Mutation characters across the 1-kb barcodes from 10x scRNA-seq cDNA libraries. **a.** Pie chart showing the frequency of different base substitution types identified in the 1-kb barcodes across all 15 individuals. C-to-T and G-to-A transitions represent 93.1% of the total editing events. **b.** Distribution of the number of cells carrying mutations at each nucleotide position along the 1-kb barcodes (HMF1 and HMF2). Each bar represents the total number of cells across all 15 individuals that harbor a mutation at that specific site.

**Fig. S8.**
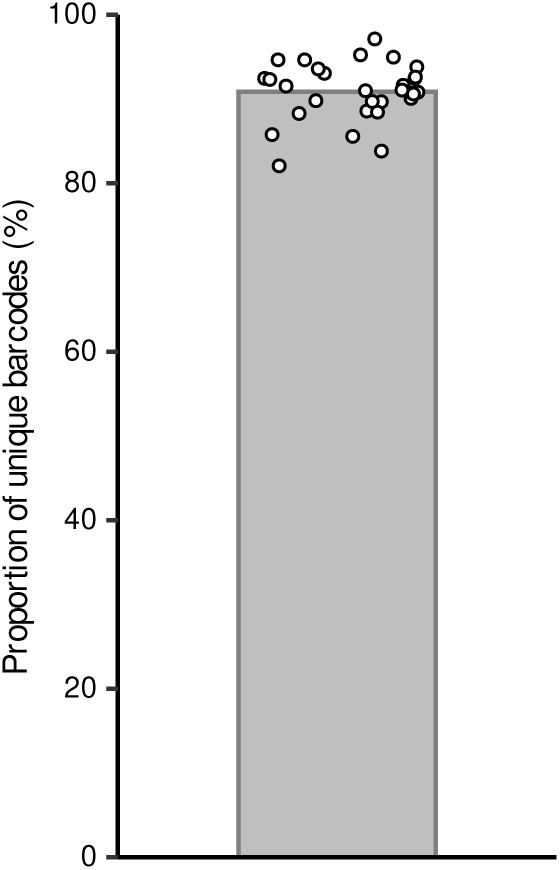
Proportion of unique 1-kb barcodes within each individual. Bar plot showing the percentage of cells carrying unique 1-kb barcodes for each HMF barcode within each individual. The bar height represents the mean proportion across all HMF barcodes (90.9%). Individual data points correspond to values calculated separately for HMF1 and HMF2 in each of the 15 individuals, yielding 30 data points in total.

**Fig. S9.**
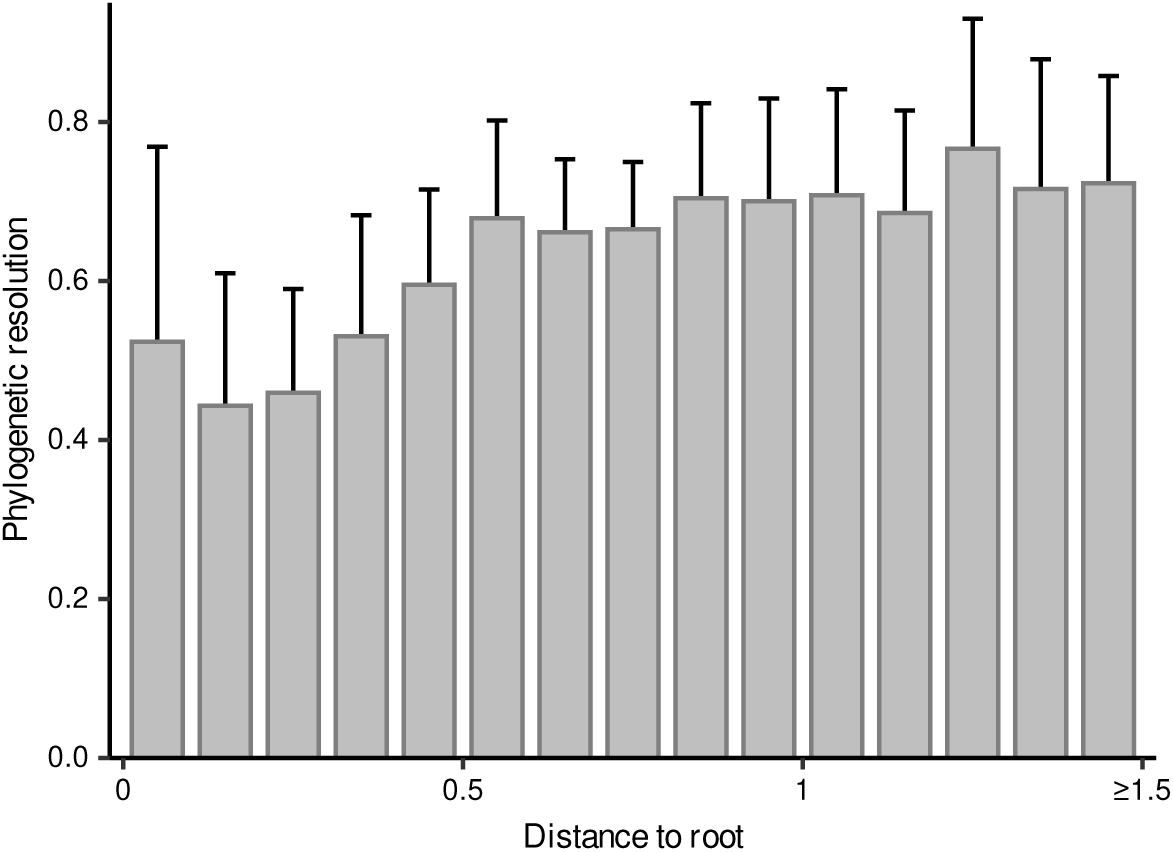
Relationship between phylogenetic resolution and the distance to root of internal nodes. Bar plot showing the phylogenetic resolution at varying tree depths across 15 individuals (n = 30 phylogenies). The x-axis represents the distance to the root, and the y-axis indicates the proportion of informative nodes within the subtrees truncated at each corresponding depth. Bars represent the mean values calculated across all 30 phylogenies, with error bars indicating the standard deviation.

**Fig. S10.**
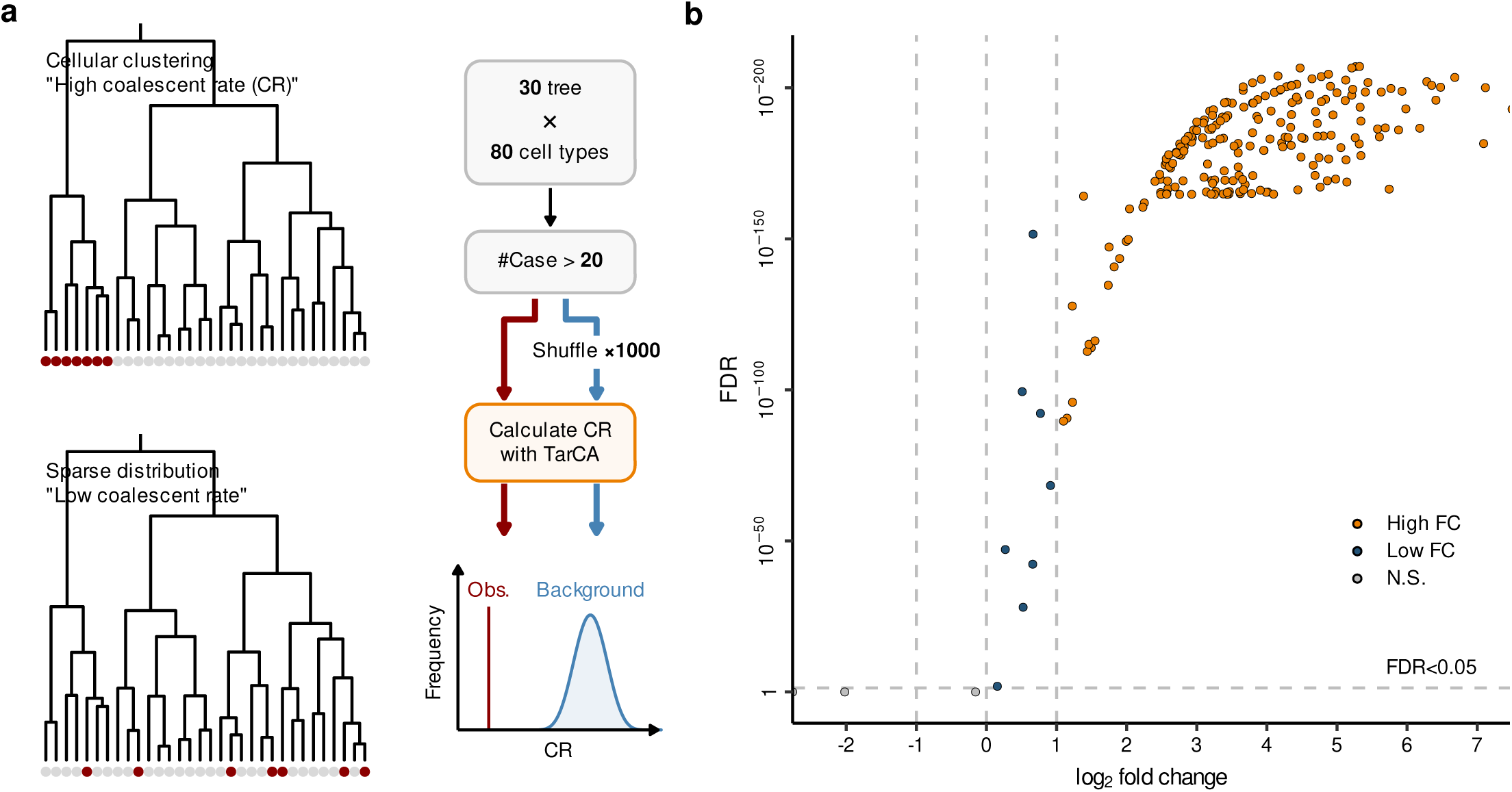
Assessment of phylogenetic co-localization for individual cell types. **a.** Schematic representation of the analysis workflow. Left: illustrative diagrams showing two scenarios of cell type distribution on a phylogeny. The top indicates cells from the same cell type clustering within a specific clade, resulting in a high coalescent rate (CR), while the bottom indicates a sparse distribution resulting in a low CR. Right: workflow for calculating phylogenetic co-localization. Observed CRs were calculated using TarCA and compared against a background distribution generated by 1,000 random shuffles of cell labels. **b.** Volcano plot showing the enrichment of CRs compared to the shuffled background. Each point represents a specific cell type within an individual phylogeny. Orange points (High FC) indicate significant phylogenetic co-localization (FDR < 0.05 and log2 fold change > 1). Dark blue points (Low FC) indicate significant results with lower effect sizes (FDR < 0.05 and log2 fold change < 1). Grey points (N.S.) represent cell types that did not reach statistical significance (FDR >= 0.05).

**Fig. S11.**
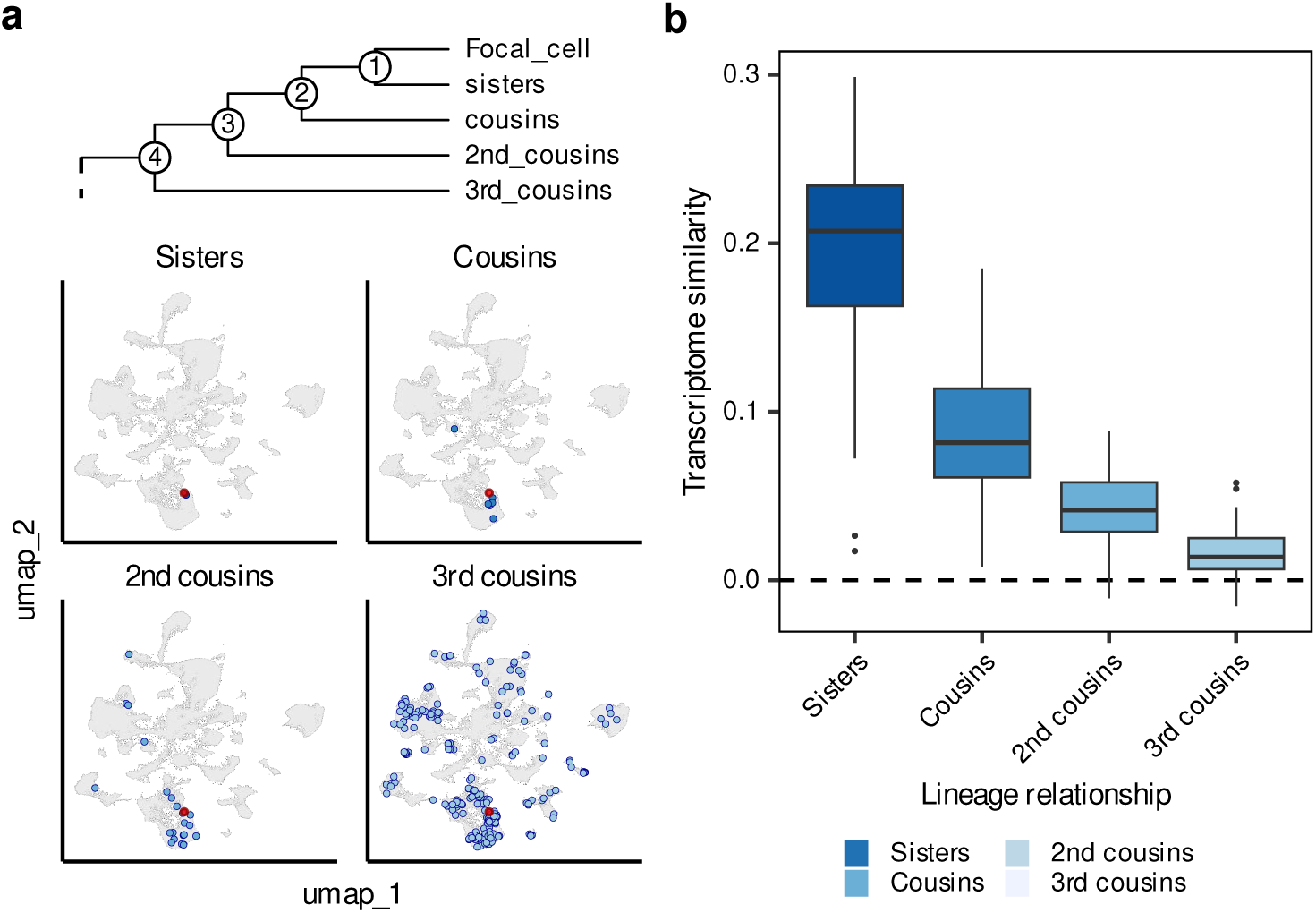
Transcriptomic similarity aligns with lineage relatedness. **a.** Illustration of cell-to-cell lineage relationships and their UMAP embeddings. Top: schematic tree showing degrees of kinship relative to a focal cell, including sisters, cousins, 2nd cousins, and 3rd cousins. Bottom: UMAP plots showing the embeddings of cells with these specific lineage relationships. The focal cell is highlighted in red, and its corresponding relatives are highlighted in blue against the background of all captured cells (grey). **b.** Boxplots showing transcriptomic similarity between cell pairs categorized by their lineage relationship. The y-axis represents transcriptomic similarity, and the x-axis categorizes cell pairs into sisters, cousins, 2nd cousins, and 3rd cousins based on their relationship in the reconstructed phylogenies. Each box represents the distribution of similarity scores for that kinship category.

**Fig. S12.**
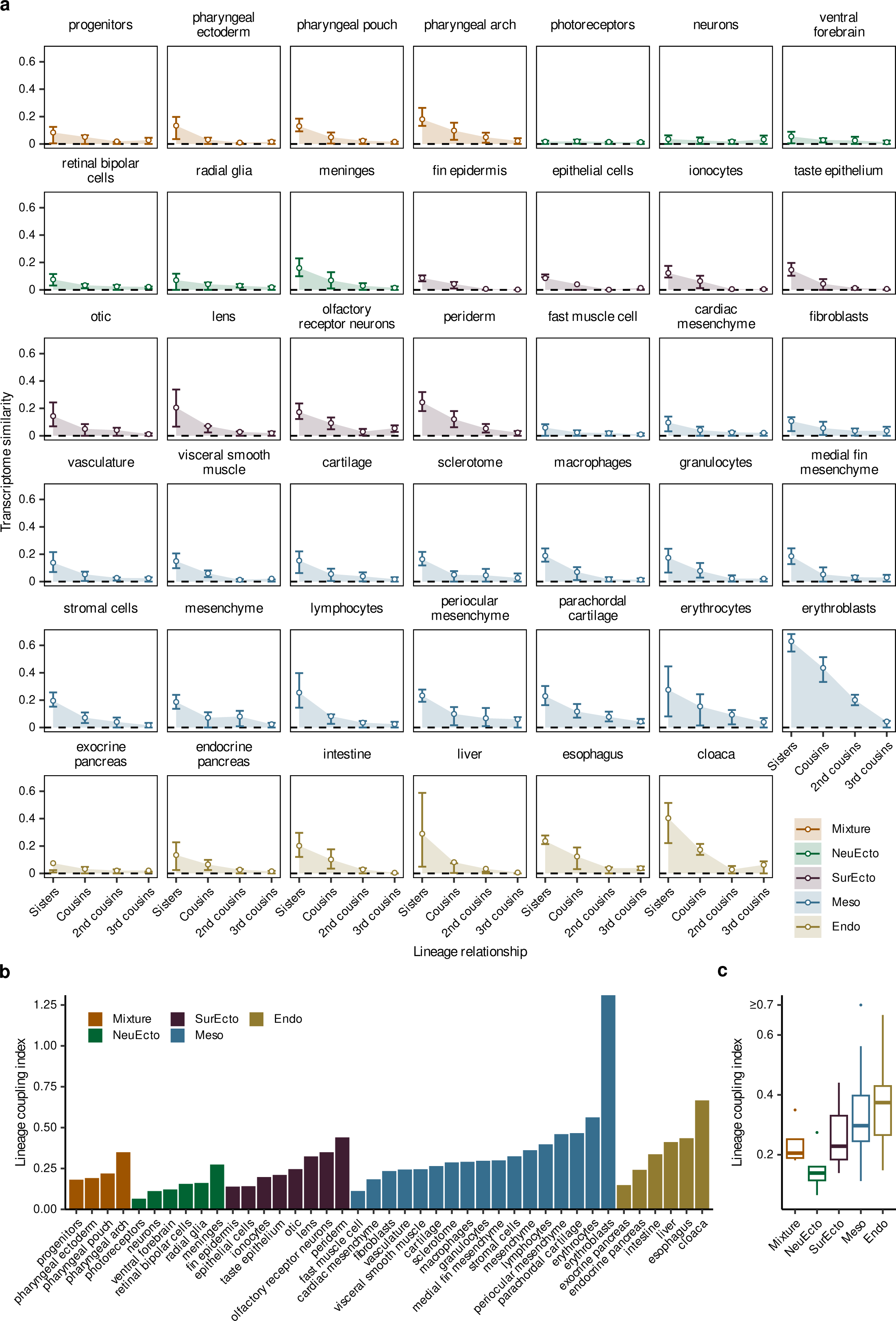
Cell-type-specific patterns of transcriptomic similarity. **a.** Transcriptomic similarity between cell pairs categorized by their lineage relationship across individual cell types. For each indicated cell type, the y-axis represents transcriptomic similarity, and the x-axis categorizes cell pairs into sisters, cousins, 2nd cousins, and 3rd cousins based on their relationship in the reconstructed phylogenies. Line colors and shaded areas correspond to the germ layer in the legend (Mixture, NeuEcto, SurEcto, Meso, and Endo). **b.** Bar plot showing the lineage coupling index for each cell type, categorized by germ layer. The lineage coupling index quantifies the strength of the relationship between transcriptomic similarity and lineage proximity. **c.** Distribution of the lineage coupling index across different germ layers. Boxplots illustrate the variation of lineage coupling within each germ layer.

**Fig. S13.**
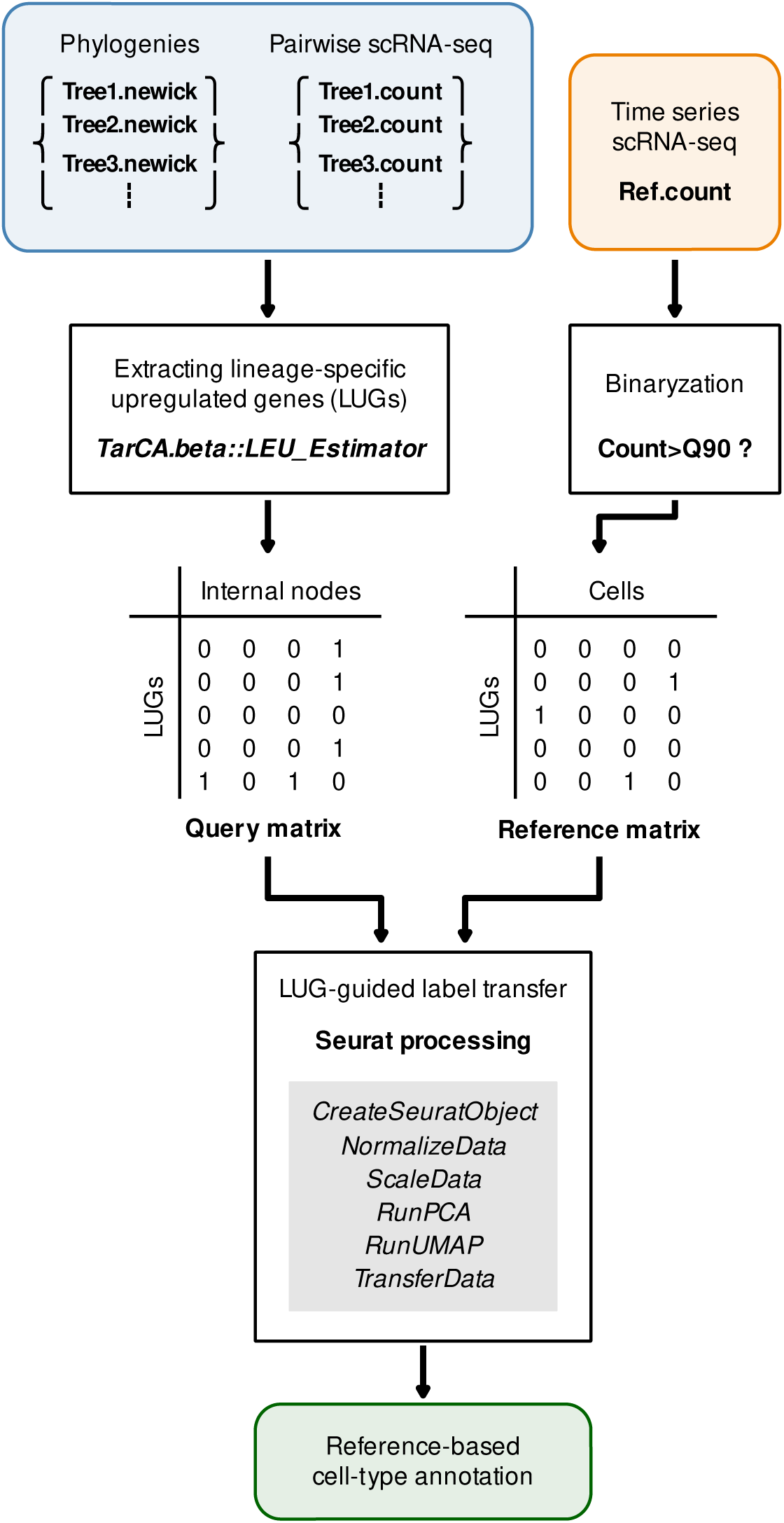
Schematic of the LEAP computational framework. Flowchart illustrating the LUG-encoded Ancestral Projection (LEAP) workflow for reconstructing ancestral transcriptional states. The process involves identifying LUG from reconstructed phylogenies and paired scRNA-seq data to generate a query matrix for internal nodes. A binary reference matrix is simultaneously constructed from time-series scRNA-seq data. These matrices are integrated via Seurat-based label transfer to project cell-type annotations from the reference onto the ancestral internal nodes.

**Fig. S14.**
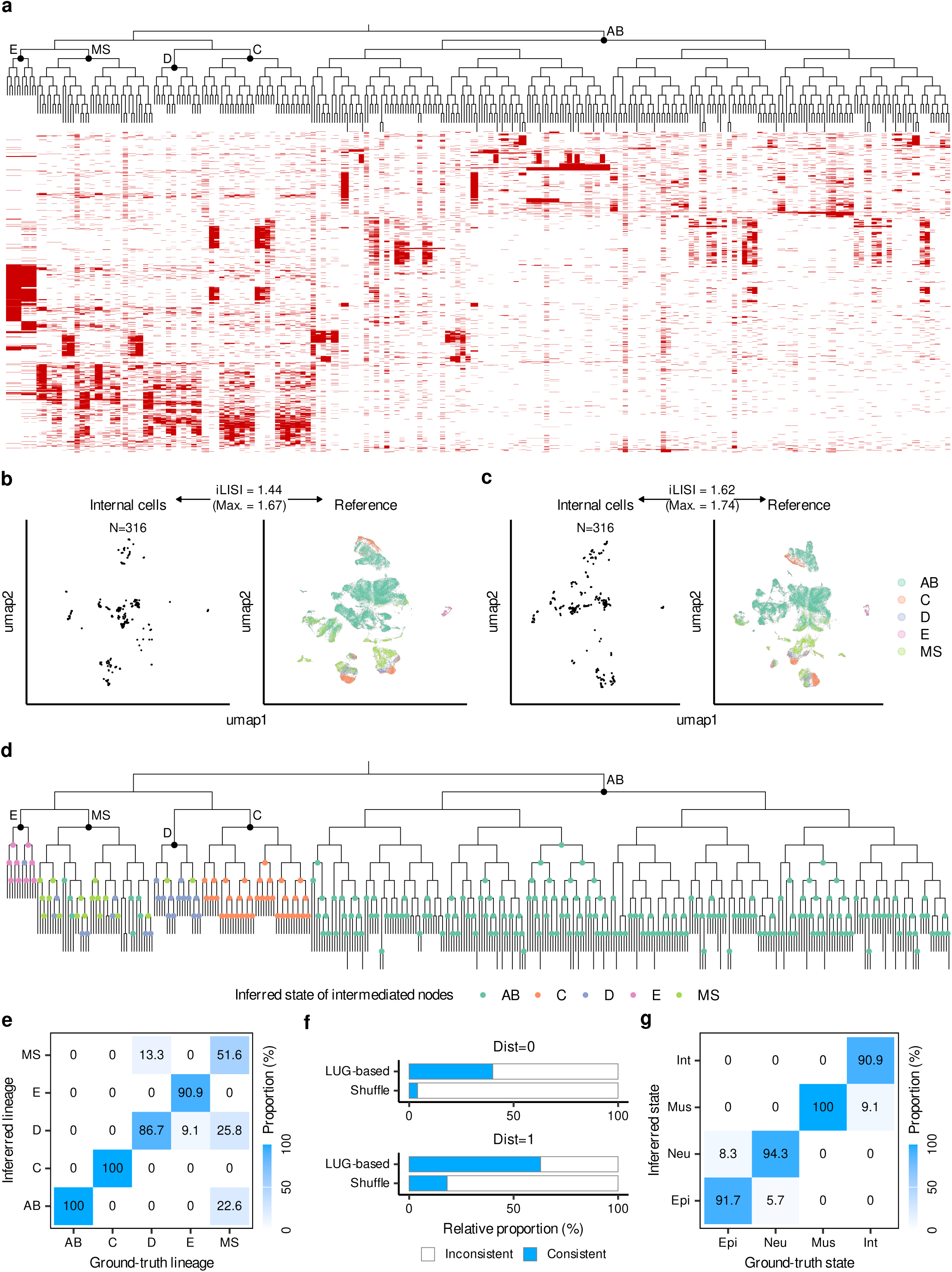
LEAP-based ancestral state inference in *C. elegans*. **a.** Heatmap showing the activation patterns of the 2,000 LUGs across the *C. elegans* phylogeny. Each red tile represents a LUG upregulated in the corresponding phylogenetic clade. **b-c.** UMAP co-embedding of reconstructed ancestral profiles (Internal cells, N=316) and the time-resolved reference atlas (N=47,798). Integration results are shown before (panel b) and after (panel c) correction, with integration Local Inverse Simpson’s Index (iLISI) values indicating the degree of mixing between datasets. Colors in the reference atlas correspond to five founder lineages (AB, C, D, E, and MS). **d.** Projection of inferred ancestral states onto the C. elegans phylogeny. Internal nodes are colored according to their predicted founder lineage assignment based on LUG-guided label transfer. **e.** Confusion matrix showing the accuracy of founder lineage assignments. The x-axis represents the ground-truth lineage, and the y-axis represents the LEAP-inferred lineage. Numbers in cells indicate the proportion (%) of nodes that were correctly or incorrectly assigned. **f.** Accuracy of identity assignment at single-cell resolution. Bar plots show the relative proportion of consistent (blue) versus inconsistent (white) assignments compared to shuffled backgrounds. "Dist=0" represents exact identity matches, and "Dist=1" represents matches within one generational distance. **g.** Confusion matrix showing the accuracy of cell-type category assignments. Reconstructed ancestral states were compared against ground-truth developmental states across four categories: Epidermis (Epi), Neurons (Neu), Muscle (Mus), and Intestine (Int).

**Fig. S15.**
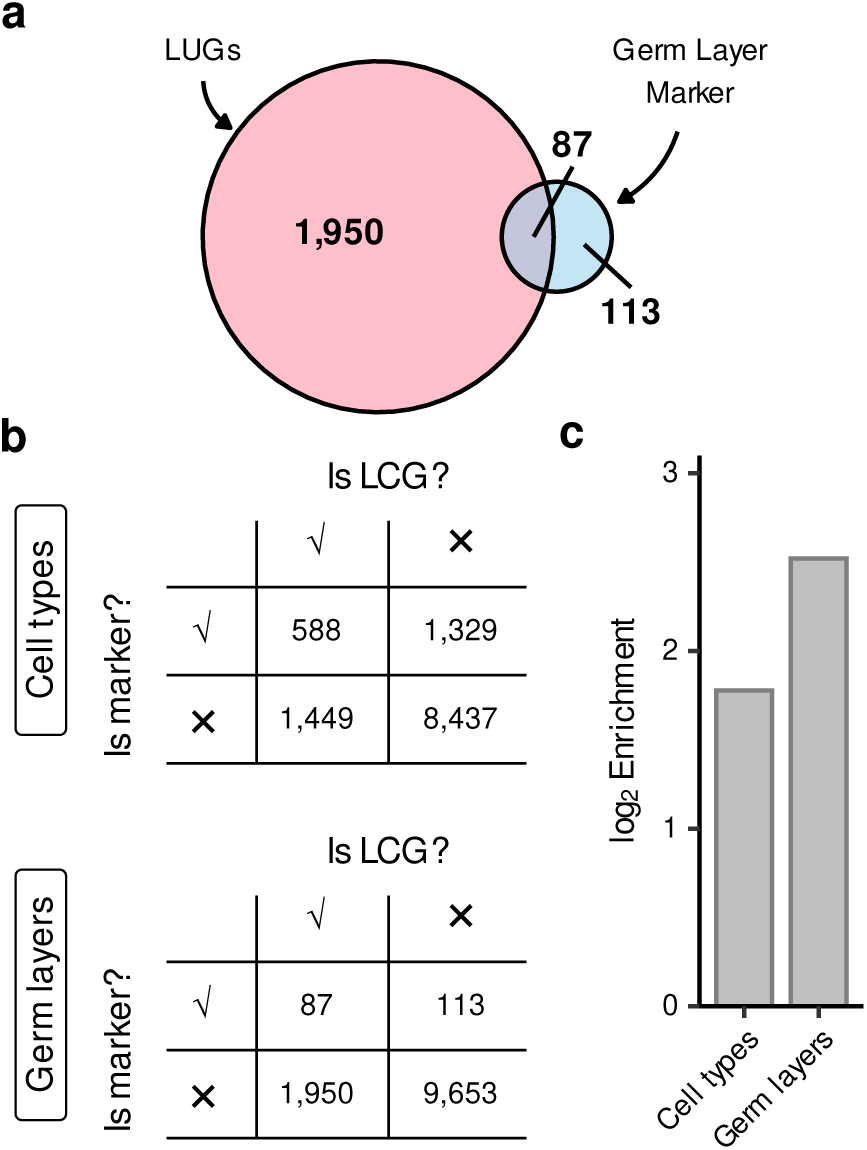
Relationship between LUGs and transcriptional markers. **a.** Venn diagram showing the overlap between LUGs and germ layer markers in zebrafish. **b.** Contingency tables evaluating the overlap between LUGs and transcriptomic markers at both the cell-type and germ-layer levels. **c.** Enrichment of transcriptomic markers within the LUG set. The bar plot shows the log2 enrichment of cell-type and germ-layer markers.

**Fig. S16.**
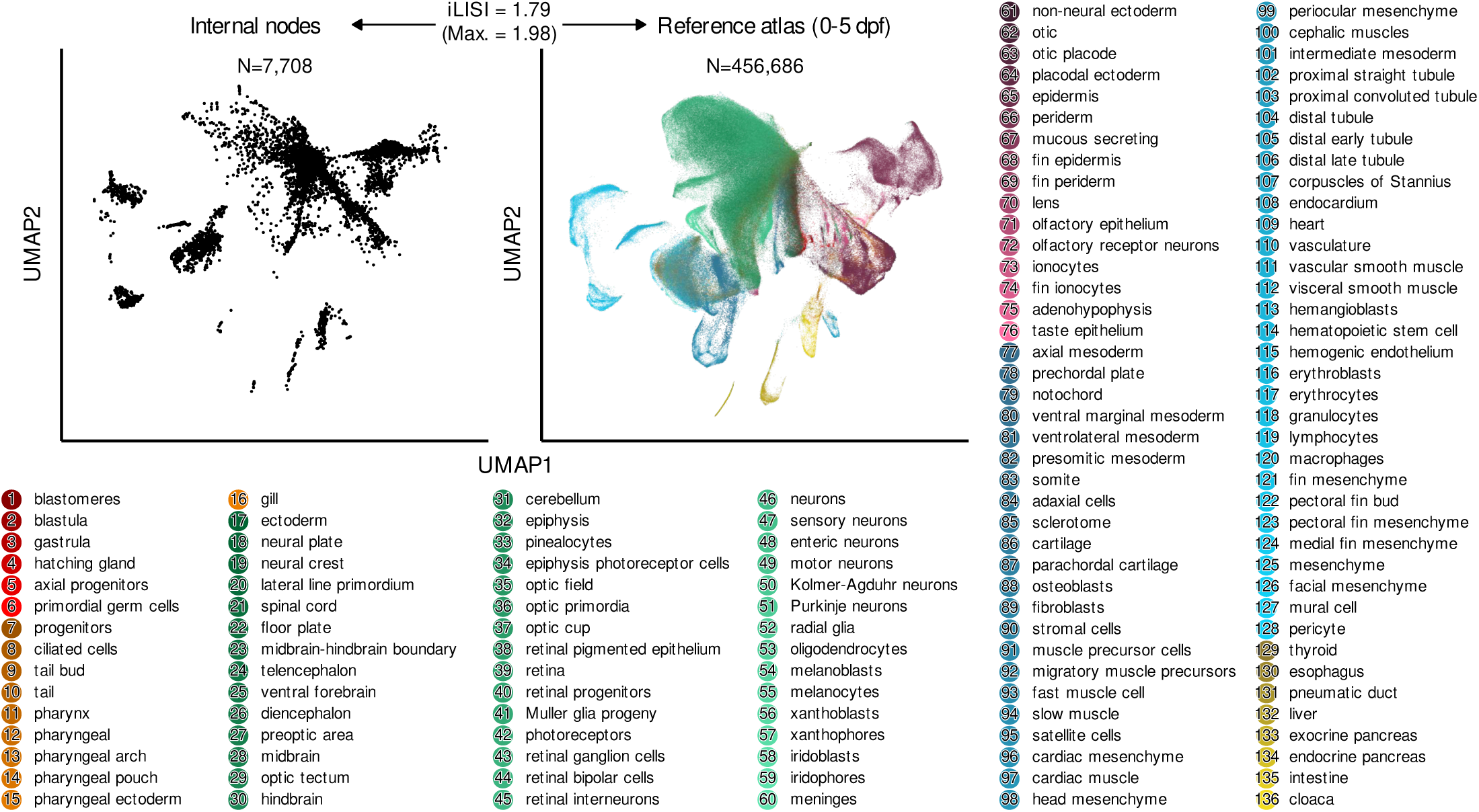
Consistency between internal profiles and reference atlas via direct-join co-embedding. UMAP co-embedding of reconstructed ancestral states (internal nodes, N = 7,708) and the time-series reference atlas (reference cells, dpf 0–5, N = 456,686). The visualization is generated via direct-join co-embedding before batch correction. Reconstructed ancestral internal nodes and reference cells are projected into a shared transcriptomic space. The integration Local Inverse Simpson’s Index (iLISI) value is 1.79 (empirical maximum = 1.98). The right panel displays the reference cells colored by their corresponding 136 cell type annotations, as detailed in the legend below.

**Fig. S17.**
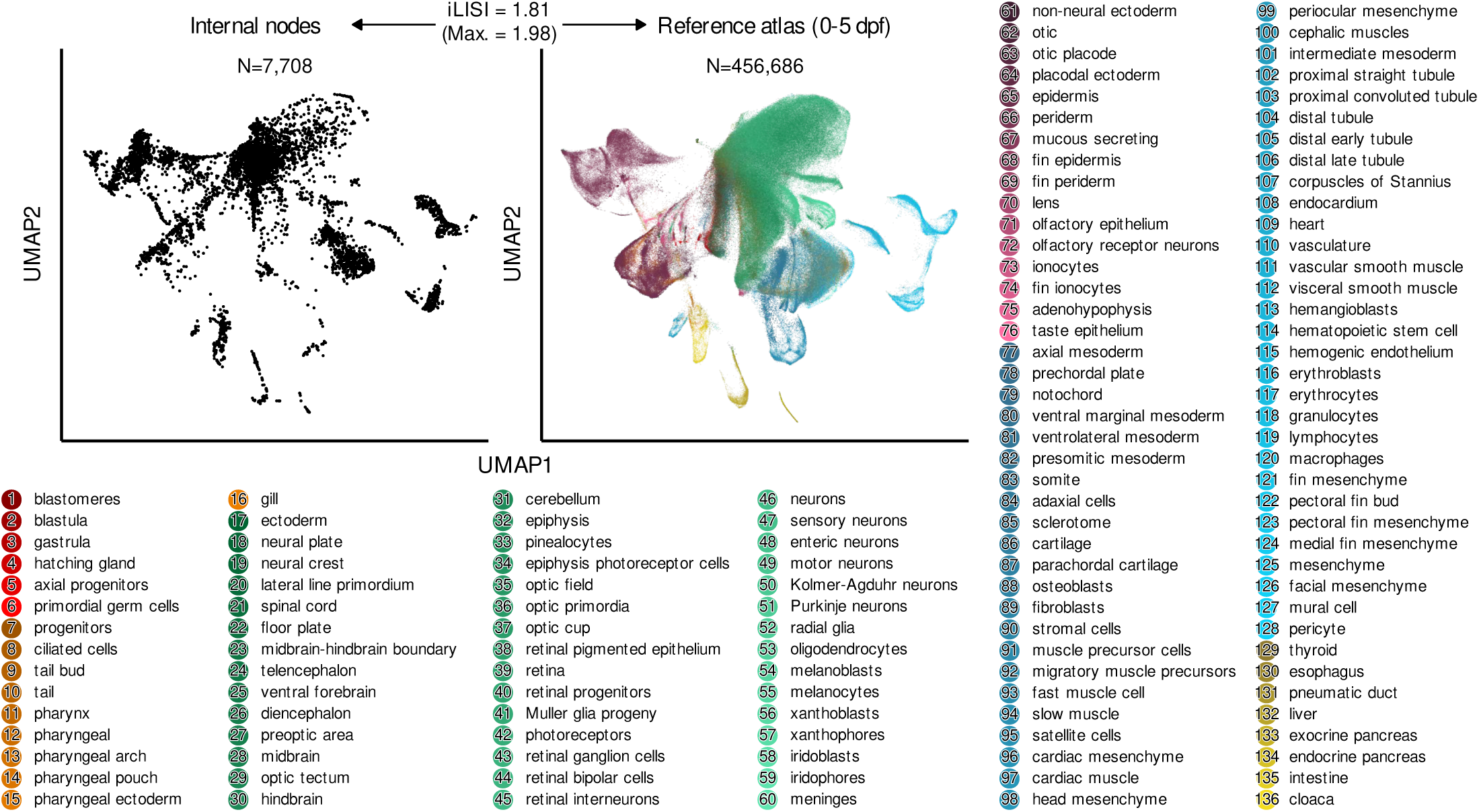
Consistency between internal profiles and reference atlas via Harmony-based co-embedding. UMAP co-embedding after Harmony integration of reconstructed ancestral states of internal nodes and the time-series reference atlas.

**Fig. S18.**
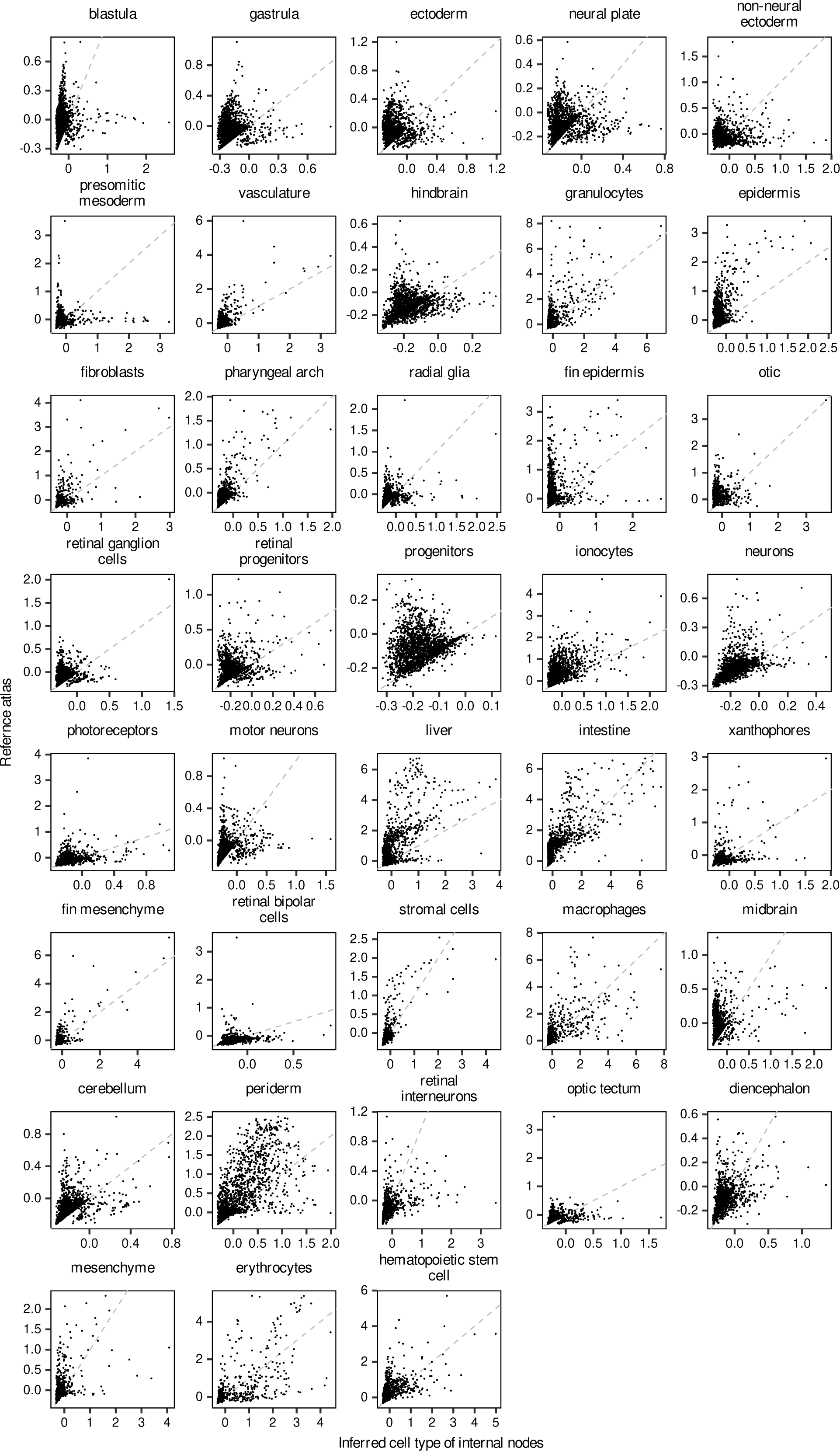
Gene expression concordance between assigned internal nodes and reference atlas. Scatter plots showing gene expression concordance between assigned internal nodes and reference cells with the same assigned cell type for the remaining 38 of the 39 cell types with more than 20 assigned internal nodes. The other cell type is shown in Fig. 4d. Each panel represents one cell type, and each point represents one gene. The x-axis indicates gene expression in internal nodes, and the y-axis indicates gene expression in reference cells. The dashed diagonal line indicates equal expression between internal nodes and reference cells.

**Fig. S19.**
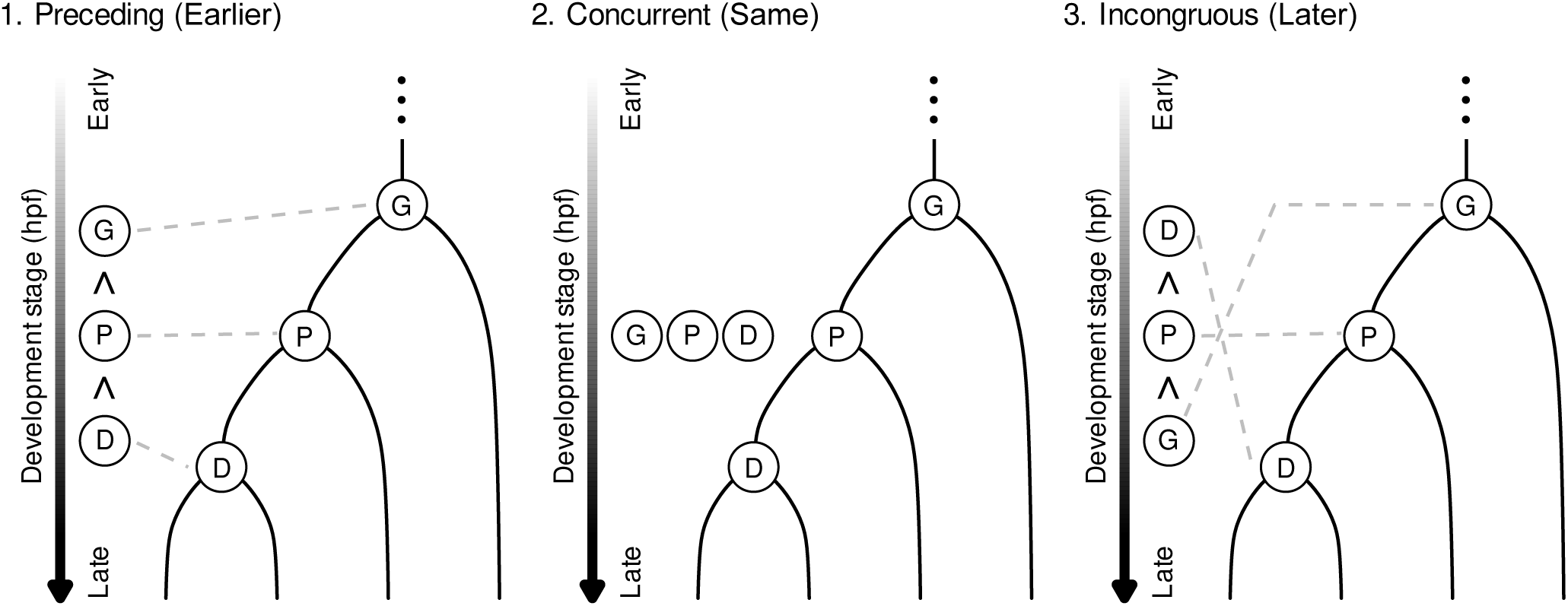
Schematic representation of temporal ordering across lineage generations. Three patterns of cell state progression from Grandmother (G) to Parent (P) and Daughter (D) stages are shown, including preceding (Earlier), concurrent (Same) and incongruous (Later).

**Fig. S20.**
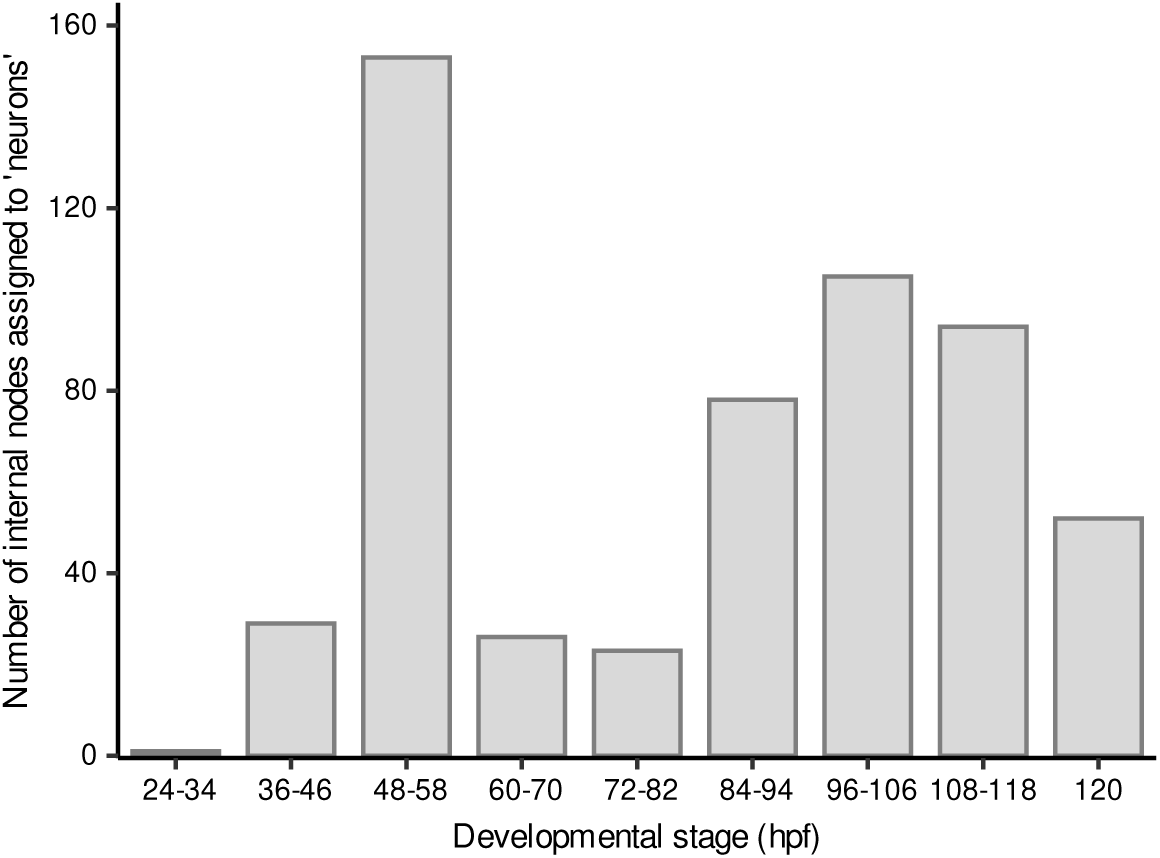
Number of internal nodes assigned to ‘neurons’ across developmental stages. Bar plot showing the number of internal nodes assigned by LEAP to the ‘neurons’ state across developmental stages. The x-axis indicates developmental stage windows in hours post-fertilization (hpf), and the y-axis indicates the number of internal nodes assigned to ‘neurons’.

**Fig. S21.**
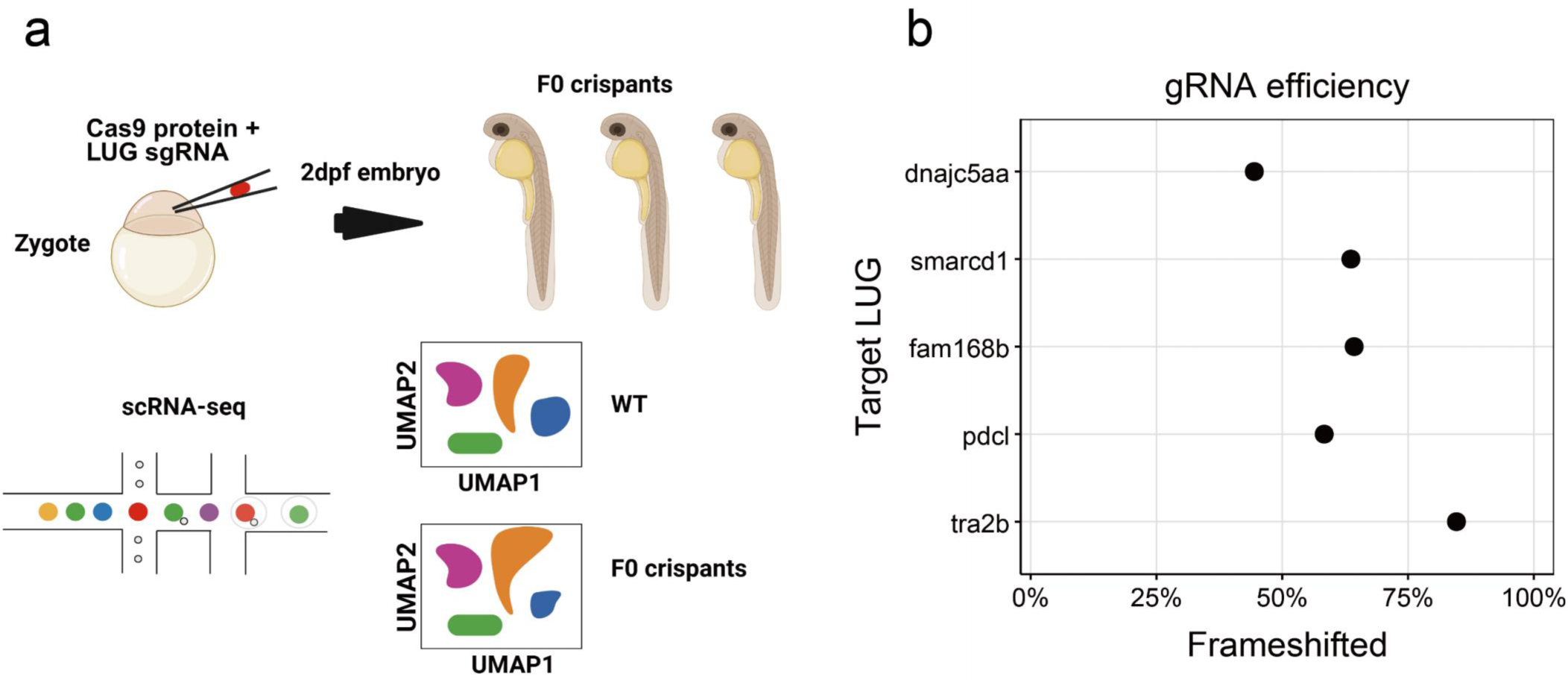
CRISPR–Cas9-mediated perturbation of candidate LUGs in zebrafish F0 crispants. **a.** Schematic overview of the F0 crispant experiment. For each selected LUG, one LUG-targeting sgRNA was injected together with Cas9 protein into one-cell-stage zebrafish embryos. WT and F0 crispant embryos were collected at 2 dpf for scRNA-seq analysis. **b.** Estimated mutagenesis efficiency of sgRNAs targeting selected LUGs based on Sanger sequencing of cloned genomic PCR products from injected F0 embryos. The x-axis indicates the percentage of frameshifted clones, and the y-axis indicates the targeted LUG.

**Fig. S22.**
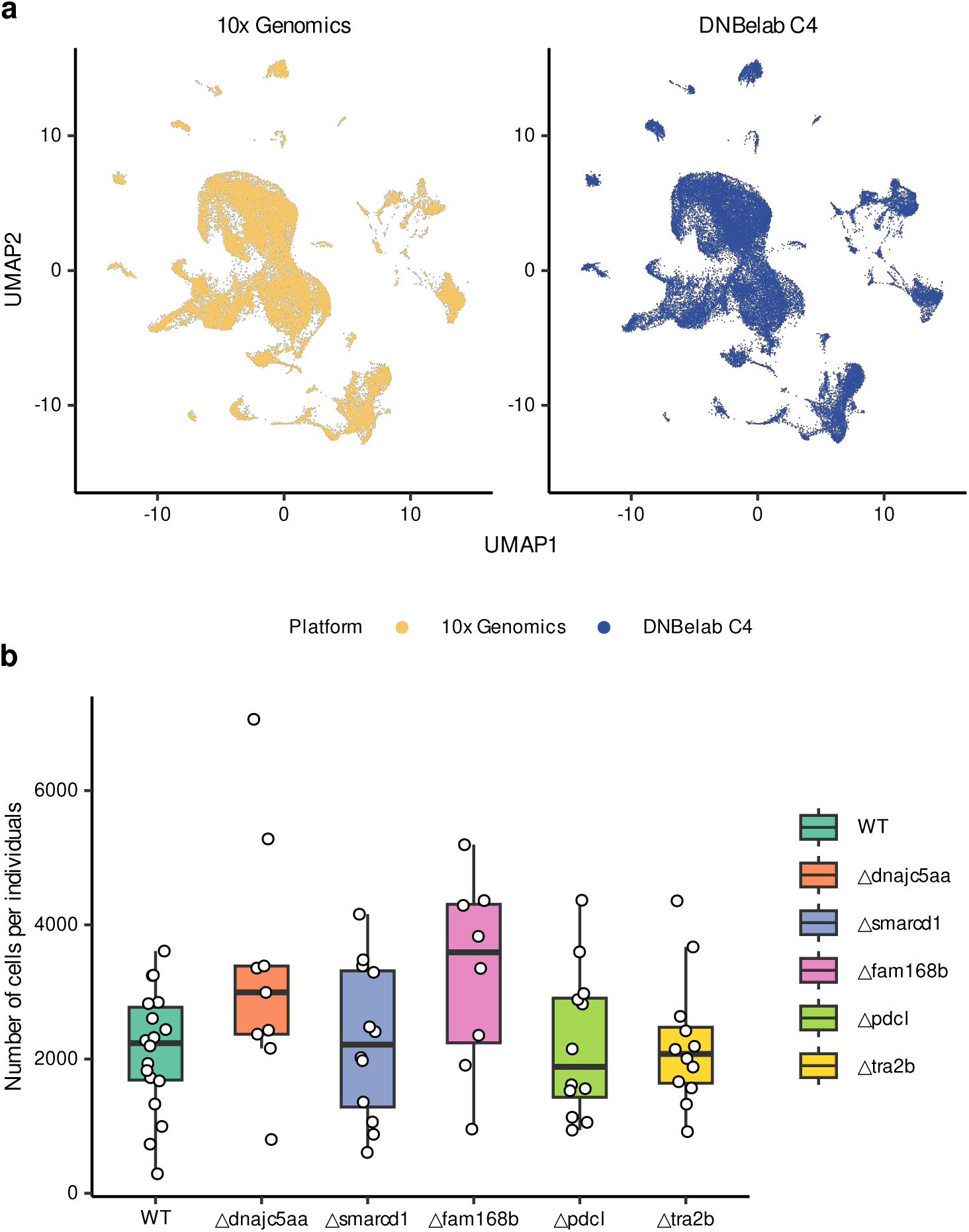
Technical comparability and sample recovery of DNBelab C4 datasets. **a.** UMAP visualization showing the global landscape of cells from 2 dpf WT embryos profiled using the 10x Genomics Chromium system (left) and the DNBelab C4 platform (right). **b.** Boxplots showing the number of cells captured per individual across wild-type (WT) and five genetic perturbation groups (dnajc5aa, smarcd1, fam168b, pdcl, and tra2b). Each data point represents a single individual larva at 2 dpf. The y-axis indicates the total number of cells captured per individual in the C4 scRNA-seq dataset. relative to WT. Bar colors denote the FDR-adjusted P-value and the direction of change, with blue and orange indicating depletion and enrichment, respectively, according to the scale shown below. NeuEcto, neuroectoderm; SurEcto, surface ectoderm; Meso, mesoderm; Endo, endoderm.

**Fig. S23.**
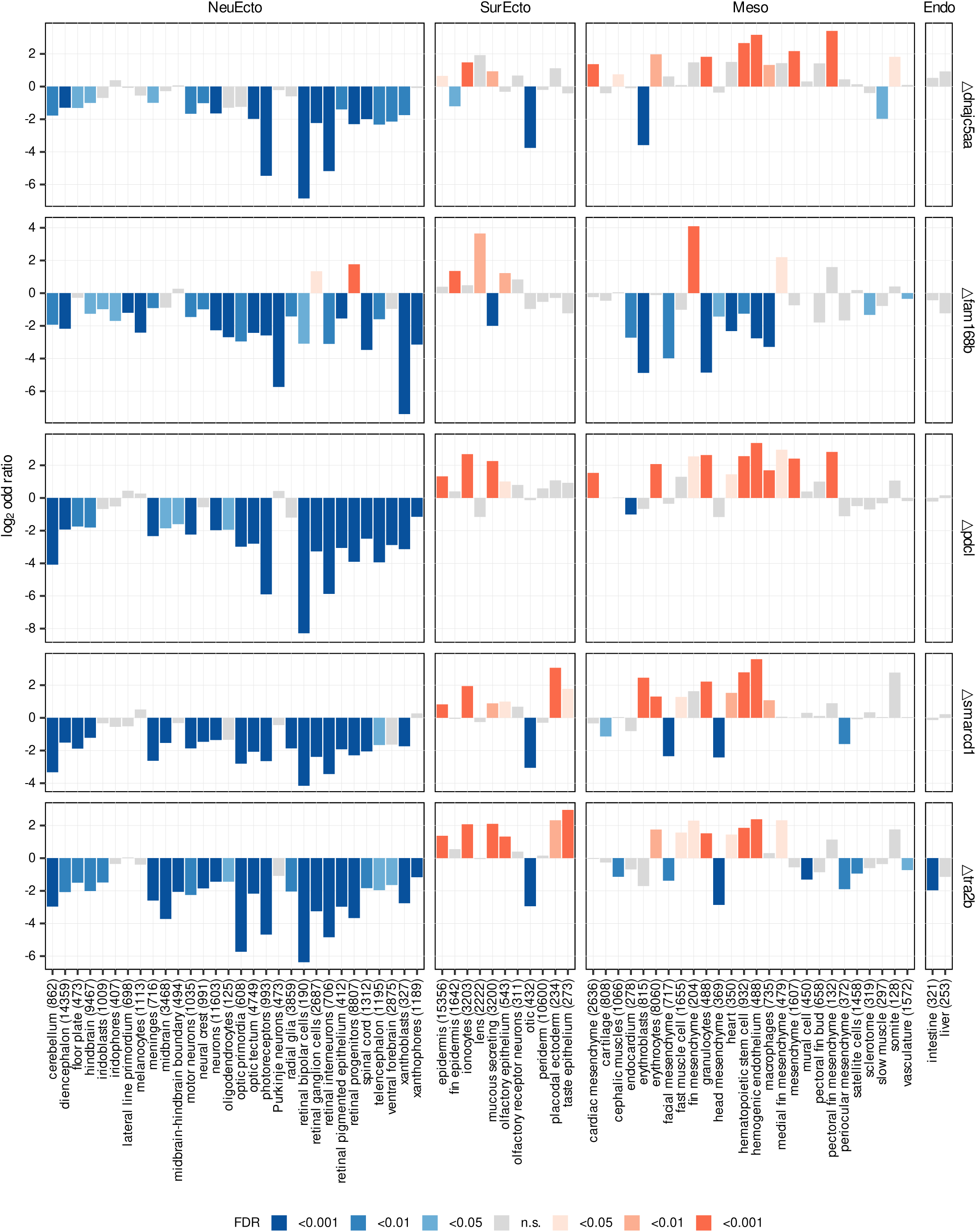
Cell-type-specific changes across five LUG perturbations. Bar plots showing changes in individual cell types for the five F0 crispant conditions shown in Fig. 5f–g. Changes are shown for each F0 crispant condition relative to the wild-type baseline. Rows correspond to the five targeted LUGs, and columns correspond to major developmental compartments (NeuEcto, SurEcto, Meso and Endo). Within each panel, each bar represents one cell type, and the numbers in parentheses on the x-axis indicate the total cell count for each type. The y-axis indicates the log2 odds ratio of the cell-type proportion in F0 crispants relative to WT. Bar colors denote the FDR-adjusted P-value and the direction of change, with blue and orange indicating depletion and enrichment, respectively, according to the scale shown below. NeuEcto, neuroectoderm; SurEcto, surface ectoderm; Meso, mesoderm; Endo, endoderm.

**Fig. S24.**
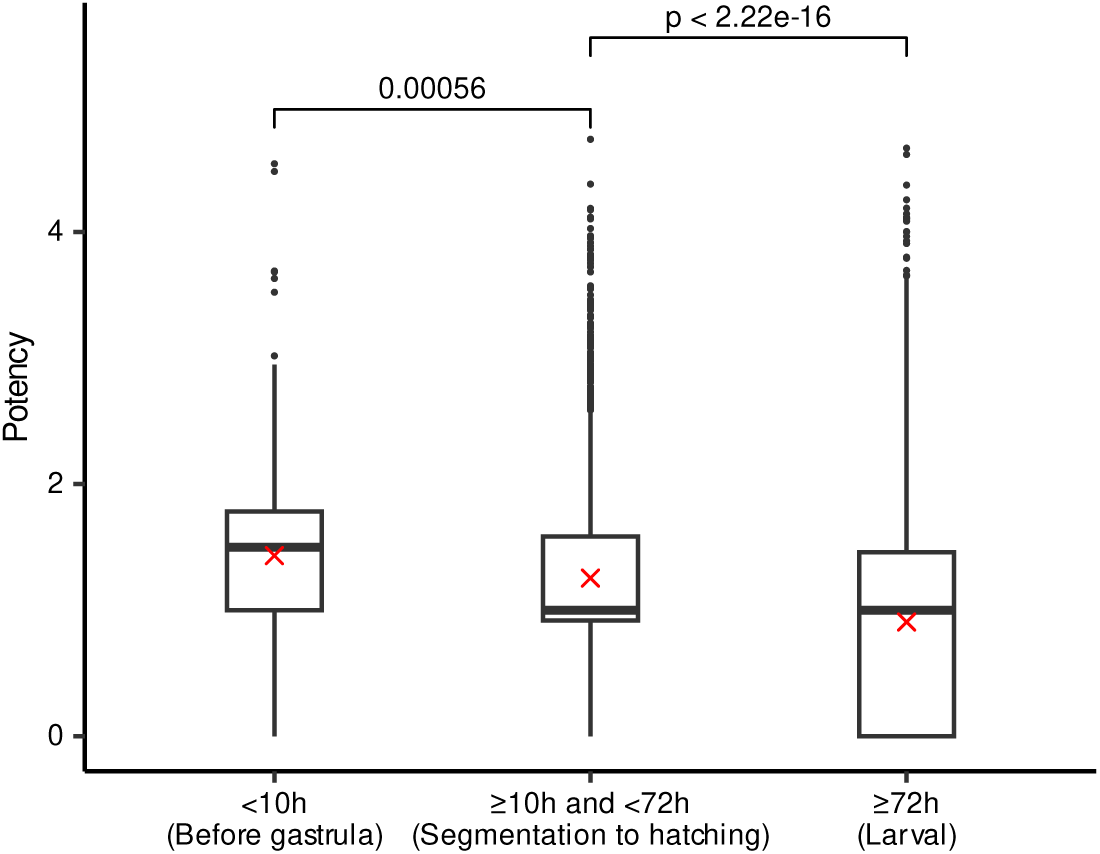
Potency across developmental stages. Boxplots showing the distribution of potency scores for internal nodes categorized into three developmental periods: early stages (up to the end of gastrulation, < 10 hpf), intermediate stages (≥10 hpf and < 72 hpf, segmentation to hatching), and late stages (≥72 hpf, larval). P-values are derived from two-sided Wilcoxon rank-sum tests.

**Fig. S25.**
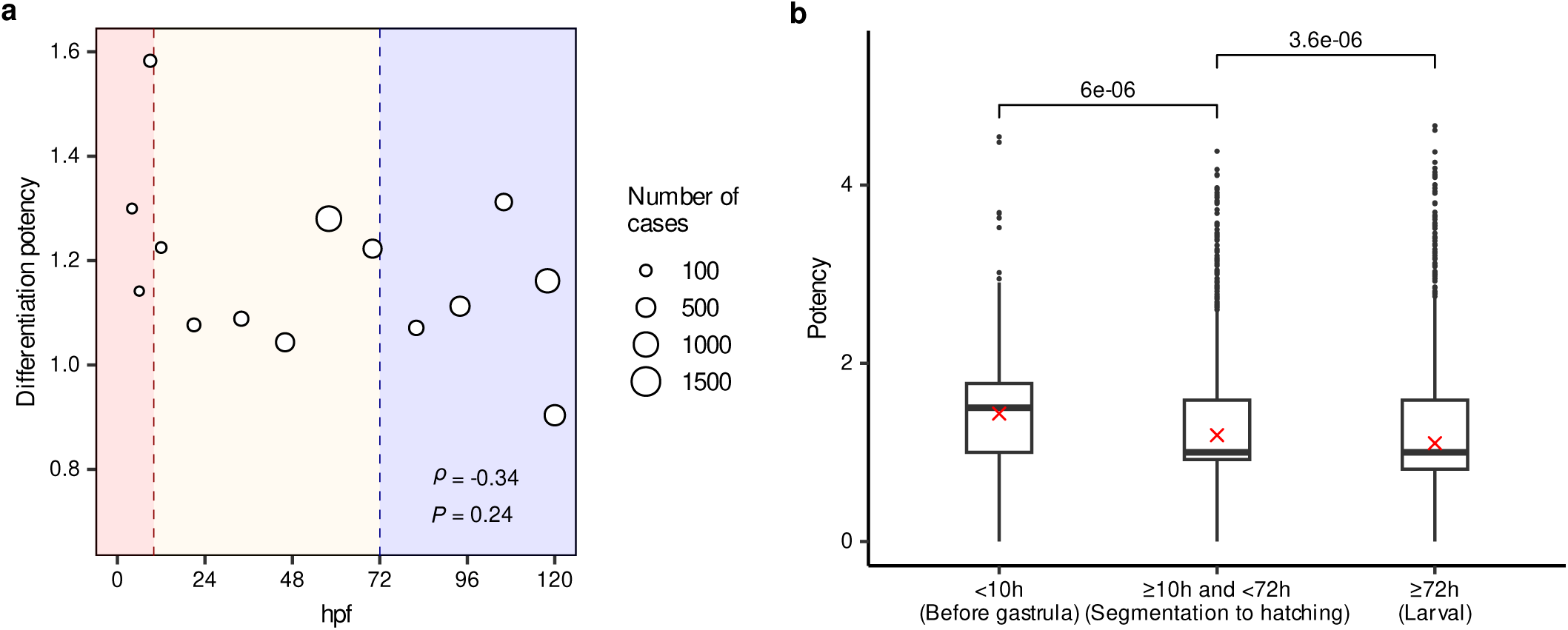
Potency across developmental stages after excluding ’periderm’ and ’progenitors’. **a.** Scatter plot showing the differentiation potency of internal nodes over developmental time (hpf), after in silico excluding cells annotated as ’periderm’ or ’progenitors’. Each circle represents a developmental stage, with the size of the circle proportional to the number of cases (internal nodes) at that stage. Spearman’s rank correlation coefficient (ρ) and the associated p-value are indicated. The background shading denotes distinct developmental periods: up to the end of gastrulation (< 10 hpf; red), segmentation to hatching (≥10 hpf and < 72 hpf; yellow), and larval stages (≥72 hpf; blue). **b.** Boxplots showing the distribution of potency scores across the three periods defined in (a), with ’periderm’ and ’progenitors’ in-silico removed from the analysis. The center line indicates the median, and the red ’x’ denotes the mean potency. P-values are calculated using two-sided Wilcoxon rank-sum tests.

**Fig. S26.**
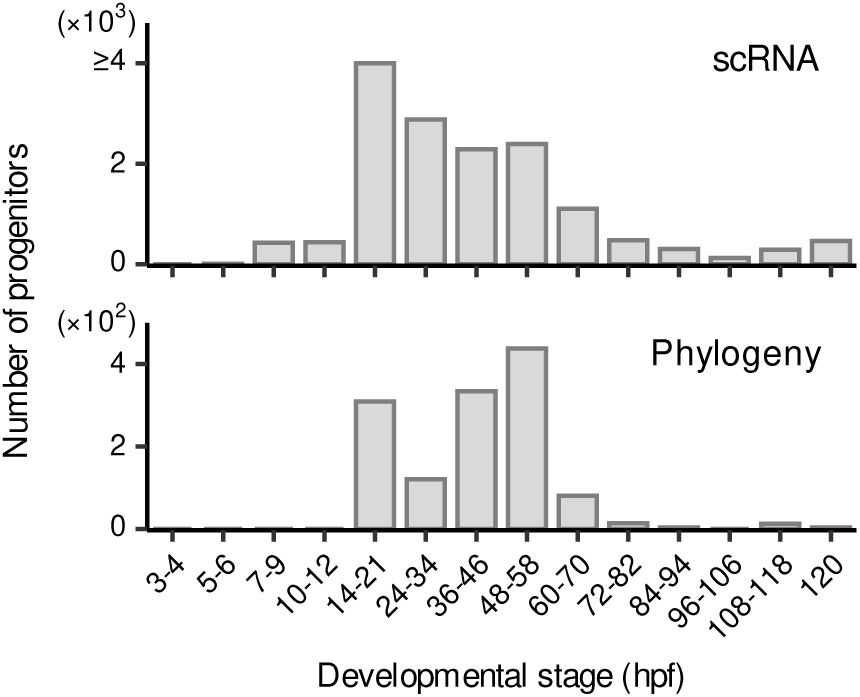
Distribution of ’progenitors’ abundance across developmental stages. Histograms showing the number of cells or internal nodes annotated as ’progenitors’ across different developmental stages (hpf). The top panel (scRNA) displays the abundance of progenitor cells annotated in the scRNA-seq reference atlas (y-axis scale: ×10^3^). The bottom panel (Phylogeny) shows the abundance of reconstructed ancestral internal nodes assigned as ’progenitors’ (y-axis scale: ×10^2^), corresponding to Fig. 6g. The x-axis represents developmental time windows in hours post-fertilization (hpf).

**Fig. S27.**
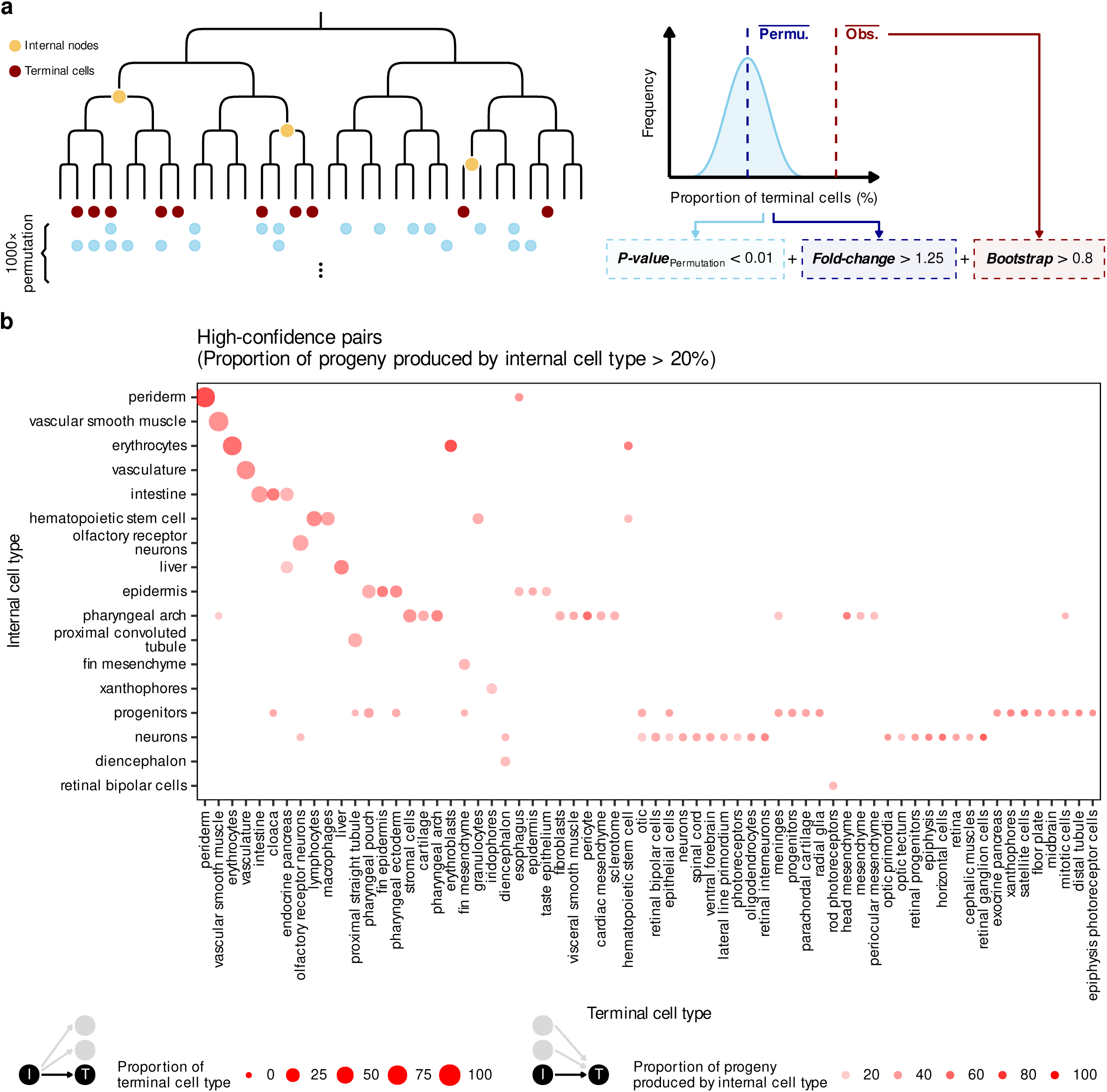
Enriched internal-to-terminal cell type associations in the cell phylogeny. **a.** Schematic overview of the filtering strategy used to select terminal cell types for phylogenetic state analysis. For each internal cell type, the proportions of its terminal descendant cell types were compared with permutation backgrounds. Internal cell type–terminal cell type pairs were retained after filtering by permutation-adjusted significance, fold change, and bootstrap support. **b.** Dot plot showing the retained internal cell type–terminal cell type pairs after filtering. The y-axis indicates internal cell types, and the x-axis indicates terminal cell types. Dot size indicates the proportion of each terminal cell type among all terminal cells, and dot color indicates the proportion of progeny from each internal cell type assigned to the corresponding terminal cell type. Terminal cell types retained in these pairs were used for subsequent phylogenetic state analysis.

**Fig. S28.**
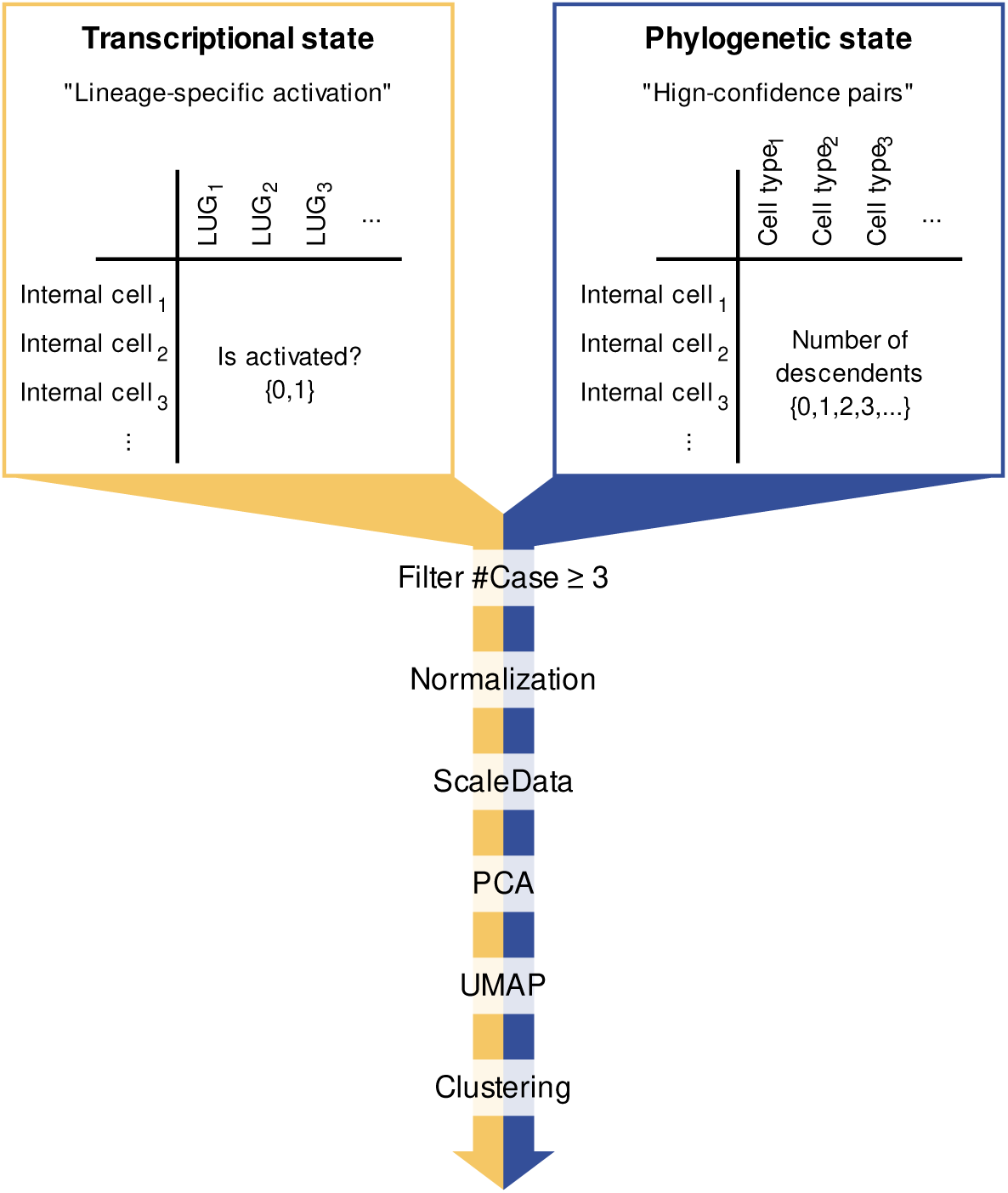
Computational workflow for characterizing phylogenetic states via terminal cell-type composition. Schematic illustration of the strategy used to cluster internal nodes according to their terminal descendant cell-type composition. For each internal node, terminal descendant cell types were summarized as a phylogenetic state matrix, in which entries represent the number of descendant cells assigned to each terminal cell type. Internal nodes with at least three high-confidence descendant cases were retained and processed using a workflow analogous to single-cell transcriptomic clustering, including normalization, scaling, principal component analysis, UMAP embedding, and clustering. The resulting clusters were used to define terminal-fate-biased subgroups within the transcriptionally similar ‘progenitors’ population.

**Fig. S29.**
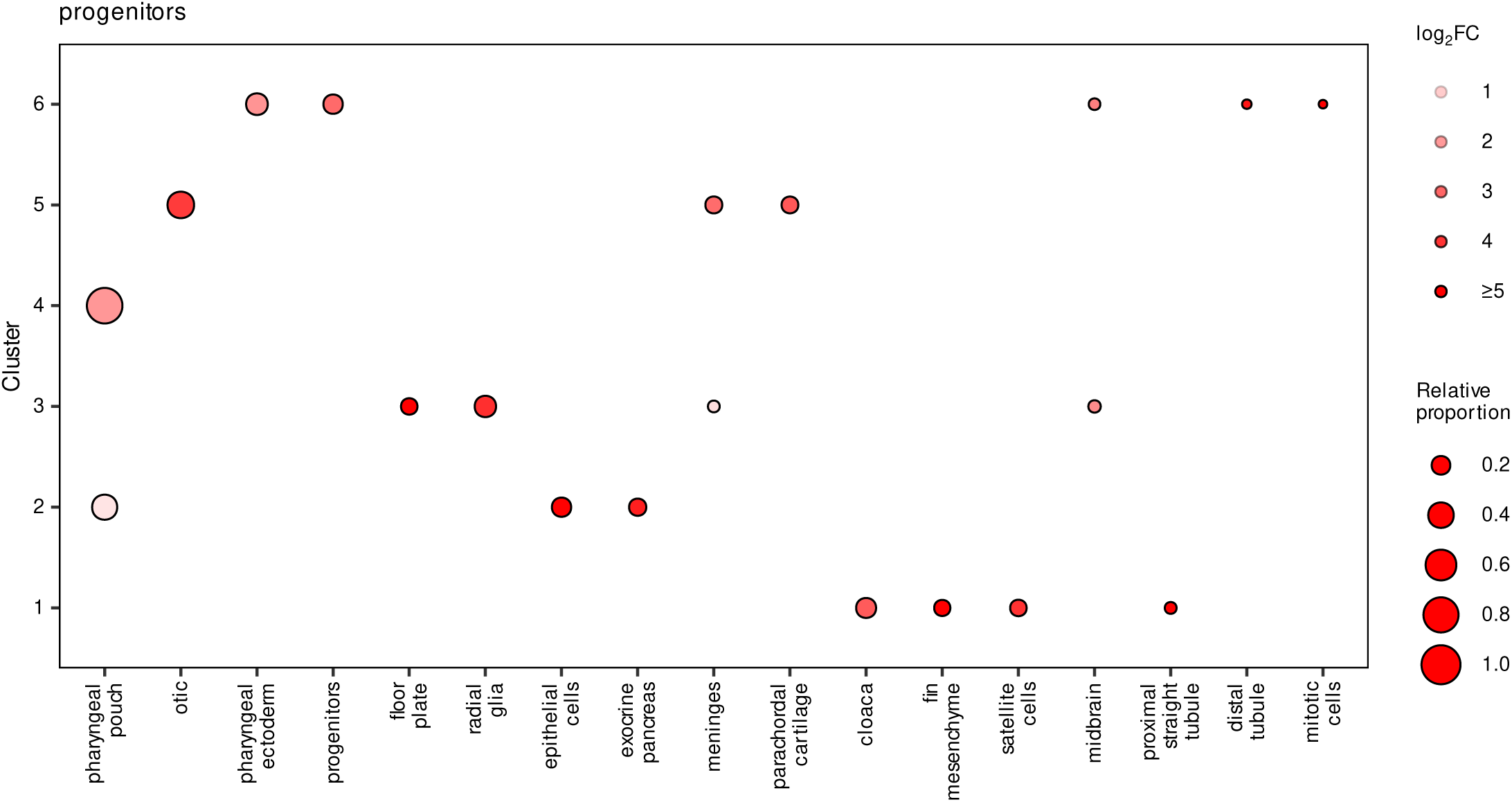
Terminal cell-type biases of phylogenetic state clusters of ’progenitors’. Dot plots showing terminal cell-type biases for phylogenetic state clusters identified with ‘progenitors’. The y-axis indicates phylogenetic state clusters, and the x-axis indicates terminal descendant cell types. Dot size indicates the relative proportion of each terminal cell type within the corresponding cluster. Dot color indicates the log_2_ fold change of the terminal cell-type proportion in that cluster relative to all other clusters.

**Fig. S30.**
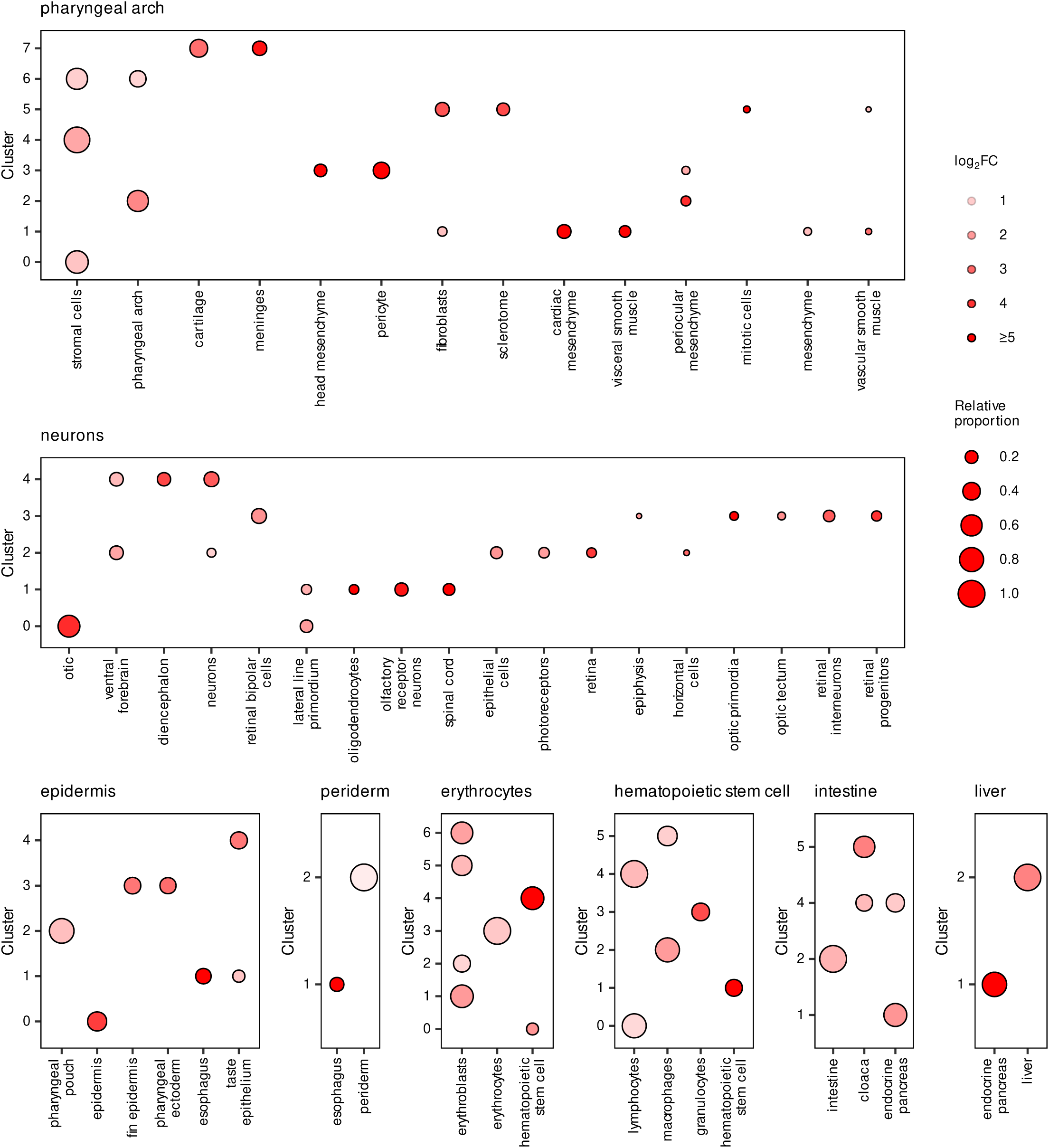
Terminal cell-type biases of phylogenetic state clusters of other cell types. Dot plots showing terminal cell-type biases for phylogenetic state clusters identified within eight internal cell types with more than 30 assigned internal nodes, including ‘epidermis’, ‘erythrocytes’, ‘hematopoietic stem cell’, ‘intestine’, ‘neurons’, ‘periderm’, ‘liver’ and ‘pharyngeal arch’. Each panel represents one internal cell type. The y-axis indicates phylogenetic state clusters, and the x-axis indicates terminal descendant cell types. Dot size indicates the relative proportion of each terminal cell type within the corresponding cluster. Dot color indicates the log_2_ fold change of the terminal cell-type proportion in that cluster relative to all other clusters.

**Supplementary Table 1 | Sequences of the 1-kb lineage barcodes HMF1 and HMF2.**

Complete sequences of the two 1-kb barcode cassettes, HMF1 and HMF2, used for lineage recording. Each barcode is 1,195 bp in length and contains a T7 promoter sequence and repeated iSceI recognition motifs. The T7 promoter, iSceI binding motif, and reverse complementary iSceI motif are highlighted as indicated in the table.

**Supplementary Table 2 | Primers used for 1-kb barcode recovery.**

List of primers used for amplification of the HMF1 and HMF2 1-kb barcode from 10x full-length cDNA library.

**Supplementary Table 3 | Estimated cell numbers and cell division numbers across zebrafish developmental stages.**

Summary of zebrafish developmental stages, corresponding developmental times, estimated cell numbers, and inferred cell division numbers. Cell numbers were obtained from published literature where available and supplemented by direct cell counting at 30 hpf in this study. The estimated division number was calculated as log₂(cell number).

**Supplementary Table 4 | Zebrafish lineage-upregulated genes identified across individual phylogenies.**

List of zebrafish lineage-specific upregulated genes (LUGs) identified from individual cell phylogenies. For each gene, the number of phylogenies in which the gene was tested and the number of phylogenies in which it was identified as significant are shown.

**Supplementary Table 5 | crRNA sequences used for CRISPR–Cas9-mediated perturbation of candidate LUGs.**

List of crRNA spacer sequences used to target the LUGs selected for CRISPR–Cas9-mediated mutagenesis in zebrafish embryos.

